# Variation in cytonuclear expression accommodation among allopolyploid plants

**DOI:** 10.1101/2022.03.10.483839

**Authors:** Corrinne E. Grover, Evan S. Forsythe, Joel Sharbrough, Emma R. Miller, Justin L. Conover, Rachael A. DeTar, Carolina Chavarro, Mark A. Arick, Daniel G. Peterson, Soraya C.M. Leal-Bertioli, Daniel B. Sloan, Jonathan F. Wendel

## Abstract

Cytonuclear coevolution is a common feature among plants, which coordinates gene expression and protein products between the nucleus and organelles. Consequently, lineage-specific differences may result in incompatibilities between the nucleus and cytoplasm in hybrid taxa. Allopolyploidy is also a common phenomenon in plant evolution. The hybrid nature of allopolyploids may result in cytonuclear incompatibilities, but the massive nuclear redundancy created during polyploidy affords additional avenues for resolving cytonuclear conflict (*i.e.,* cytonuclear accommodation). Here we evaluate expression changes in organelle-targeted nuclear genes for six allopolyploid lineages that represent four genera (*i.e., Arabidopsis, Arachis, Chenopodium*, and *Gossypium*) and encompass a range in polyploid ages. Because incompatibilities between the nucleus and cytoplasm could potentially result in biases toward the maternal homoeolog and/or maternal expression level, we evaluate patterns of homoeolog usage, expression bias, and expression level dominance in cytonuclear genes relative to the background of non-cytonuclear expression changes and to the diploid parents. Although we find subsets of cytonuclear genes in most lineages that match our expectations of maternal preference, these observations are not consistent among either allopolyploids or categories of organelle-targeted genes. Our results indicate that cytonuclear expression accommodation may be a subtle and/or variable phenomenon that does not capture the full range of mechanisms by which allopolyploid plants resolve nuclear-cytoplasmic incompatibilities.

## Introduction

Intergenomic coevolution between the nucleus and organelle(s) is a common feature among eukaryotes. Gene loss and transfers to the nucleus have greatly reduced the coding regions of modern mitochondrial and plastid genomes to a limited number of essential genes (Greiner and Bock 2013; Budar and Mireau 2018; Giannakis *et al*. 2021). Consequently, these organelles must coordinate transcripts and protein products from two or more different genomic compartments to carry out essential cellular functions. Over time, this functional interdependence results in coadaptation between the nucleus and each organelle; however, differences in mode of inheritance (i.e., biparental for the nucleus and cytoplasmic for the organelles) can lead to incompatibilities between nuclear and organellar alleles, particularly in hybrid lineages. These cytonuclear incompatibilities are widespread among species and can have dramatic consequences for fitness (Fishman and Willis 2006; Hill 2017; Fishman and Sweigart 2018; Postel and Touzet 2020), even leading to hybrid breakdown in some cases (Burke and Arnold 2001; Greiner *et al*. 2011; Burton and Barreto 2012; Burton *et al*. 2013; Budar and Mireau 2018).

Cytonuclear incompatibilities arising when evolutionarily distinct lineages merge to form allopolyploids may experience additional complex fates compared to incompatibilities in homoploid lineages (Sharbrough *et al*. 2017). The combined effects of genome merger and doubling have generally been associated with a diverse array of genomic and transcriptional changes, including nonrandom gene loss, intergenomic gene conversion, and epigenetic/regulatory changes leading to (sometimes biased) alterations in gene expression (Chen 2007; Doyle *et al*. 2008; Freeling 2009; Gaeta and Pires 2010; Jackson and Chen 2010; Salmon *et al*. 2010; Grover *et al*. 2012; Madlung and Wendel 2013; Yoo *et al*. 2014; Song and Chen 2015; Bao *et al*. 2019; Gallagher *et al*. 2020). While often evaluated on an individual gene basis, many genes are sensitive to the abundance of interacting partners, particularly those involved in multi-subunit complexes (Birchler and Veitia 2010, 2014, 2021). In allopolyploid lineages, coordination of gene products becomes more complicated when interactions between previously isolated genomes occur and redundancy affords the possibility of gene loss or divergence (Adams and Wendel 2005; Conant and Wolfe 2008; Buggs *et al*. 2011; Conant *et al*. 2014; Gout and Lynch 2015; Panchy *et al*. 2016; Cheng *et al*. 2018; Nieto Feliner *et al*. 2020).

While cytonuclear incompatibilities arising in homoploid hybrid species and their roles in homoploid hybrid speciation have been described for many species (Levin 2003; Greiner *et al*. 2011; Burton and Barreto 2012; Burton *et al*. 2013; Sloan *et al*. 2017), the problem of maintaining coordinated expression after genome merger coupled with whole genome duplication has only recently been considered and may be particularly acute for nuclear-encoded organelle-targeted proteins whose organelle-encoded interacting partners derive from only one of the two parents (Sharbrough *et al*. 2017). In addition to issues surrounding parental divergence and potential copy number variability in some organelle-interacting genes, allopolyploid species both face additional challenges relating to their massive duplication, including nucleotypic effects (Doyle and Coate 2019), and harbor additional mechanisms for resolving conflict, such as homoeologous exchange (Gaeta and Pires 2010; Bird *et al*. 2018; Mason and Wendel 2020).

Consequently, a number of co-evolutionary processes might operate to balance the interaction between the nucleus and organelles, including copy number changes in organelle-interacting nuclear genes, increased organellar biogenesis, up-regulation of maternal and/or organellar genes with concomitant paternal down-regulation, selection for gene conversion or other mutations favoring maternal-like sequences, and pseudogenization of incompatible paternal copies (Sharbrough *et al*. 2017; Doyle and Coate 2019).

Recent research has begun to shed light on the extent and consequences of cytonuclear incompatibility in polyploid species. One of the first examples came from the genus *Gossypium*, in which the Rubisco complex exhibits maternally biased homoeolog retention, expression levels, and asymmetric gene conversion (Gong *et al*. 2012), and these observations were extended for Rubisco in phylogenetically disparate allopolyploids including *Arabidopsis*, *Arachis*, *Brassica*, and *Nicotiana* (Gong *et al*. 2014). Similar results were seen for the organelle-interacting gene *MS1* in allohexaploid wheat (ABD genomes in a B cytoplasm), where only B-homoeologs exhibited expression, and homoeologs from the non-matching (AD) genomes were epigenetically silenced (Wang *et al*. 2017b). The recently formed allotetraploid *Tragopogon miscellus* also exhibited maternal bias for cytonuclear related genes, but only for a subset of the naturally occurring *T. miscellus* individuals surveyed and none of the synthetic individuals (Sehrish *et al*. 2015; Shan *et al*. 2020). Similar observations were made for synthetic allopolyploids from *Cucumis* (Zhai *et al*. 2019), rice (Wang *et al*. 2017a), and in both the recent natural and newly synthesized forms of allopolyploid *Brassica* (Ferreira de Carvalho *et al*. 2019), suggesting that cytonuclear coordination may not occur immediately in nascent polyploid species.

Here we examine the evolutionary consequences of genome merger and doubling on the expression of nuclear-encoded genes whose products are targeted to the mitochondria or plastids and interact with mitochondrial and/or plastid gene products (*i.e*., cytonuclear genes). Using five independent polyploid events in four genera that encompass a range of polyploid ages and diploid divergence times, we quantify patterns of homoeolog usage in cytonuclear genes and patterns of total expression. We look for evidence of cytonuclear accommodation by testing the hypotheses that cytonuclear genes of allopolyploid taxa exhibit (1) maternally biased homoeolog expression and/or (2) maternal expression level dominance (*i.e.,* expression patterns that more closely resemble maternal diploids than paternal diploids), reflecting a response to the historical coevolution between the maternal subgenome and the maternally inherited organelles.

## Methods

### Plant Materials and sequencing

Five plants were grown for each diploid and polyploid representative from four genera: *Arabidopsis, Arachis, Chenopodium*, and *Gossypium* (Supplementary Table 1). Growth conditions for each genus are listed below.

#### Arabidopsis

Allopolyploid *Arabidopsis suecica* (*Arabidopsis thaliana* x *Arabidopsis arenosa*) accession CS22505 seeds were acquired from Andreas Madlung (University of Puget Sound, Washington USA). These were grown in a common incubator with representatives of the parental species, *Arabidopsis arenosa* (paternal, accession CS3901xKB3) and *Arabidopsis thaliana* Landsberg *erecta* (maternal) whose seeds were provided by Roswitha Shmickl (Charles University, Prague) and Andreas Madlung, respectively. Seeds were surface sterilized using 70% v/v ethanol and placed on Murashige and Skoog (MS) plates for vernalization and germination. After the vernalization period (*i.e.*, two weeks at 4 °C), plates were moved to their growing conditions (20°C, 16/8 hours light/dark). Once germinated, seeds were moved to 6-inch diameter pots with potting soil (Sungro SUN52128CFLP). After several weeks of growth, plants were winterized (8°C, 10/14 hours light/dark) to induce flowering. Once plants were mature, leaves were harvested from each plant at a uniform time of day (midday) and flash frozen for RNA extraction.

#### Arachis

*Arachis* was represented by two allopolyploid genotypes, *i.e.*, *Arachis hypogea* cv. Tifrunner (Holbrook and Culbreath 2007) and the synthetic (*Arachis ipaensis* x *Arachis duranensis*)^4x^ known as IpaDur1 ((Fávero *et al*. 2006; Leal-Bertioli *et al*. 2018); hereafter *Arachis* IpaDur1), as well as their two model diploid progenitors, *Arachis duranensis* (accession V14167) and *Arachis ipaensis* (accession K30076). Notably, these two allopolyploid species have opposite parentage; *Arachis duranensis* is maternal for *Arachis hypogea* but paternal for *Arachis* IpaDur1. All species were grown in an environmentally-controlled greenhouse at the University of Georgia. The first expanded leaves were collected from eight-week-old plants; these were flash frozen in liquid nitrogen and shipped on dry ice to Iowa State University for RNA extraction.

#### Chenopodium

The allopolyploid species *Chenopodium quinoa* accession QQ74 was grown along with the model progenitor species *Chenopodium pallidicaule* (maternal; PI 478407) and *Chenopodium suecicum* (paternal) by David Brenner in the United States Department of Agriculture (USDA, Ames, Iowa) greenhouse at Iowa State University and provided as living material. Samples were harvested directly from the greenhouse at a uniform time of day and flash frozen in liquid nitrogen for RNA extraction.

#### Gossypium

*Gossypium* was represented by two allopolyploid species, *i.e.*, *Gossypium hirsutum* cultivar TM1 and *Gossypium barbadense* accession GB379, and their two model diploid progenitors, *Gossypium arboreum* (maternal) and *Gossypium raimondii* (paternal). Samples were grown from seed in a common environment in the Pohl Conservatory at Iowa State University.

Seeds were planted in 2 gallon pots with a custom potting mixture of 4:2:2:1 Sungro soil : perlite : bark : chicken grit. *Gossypium* was grown to maturity (minimum of 6 months) under typical greenhouse conditions, collected at a uniform time of day, and flash frozen in liquid nitrogen for RNA extraction.

#### All plants

A minimum of five replicates (leaf tissue) were collected for each species. RNA was extracted from the *Arabidopsis*, *Arachis*, and *Chenopodium* samples using the Direct-zol RNA kit (Zymo Research), including 600ul of Trizol. For *Arachis,* an additional grind step in 600ul of Trizol using ⅛ inch diameter steel beads (1-2 minutes of vortexing) immediately followed the initial grind in liquid nitrogen, and 400ul of additional Trizol was added for extraction. All other steps follow the manufacturer protocol. *Gossypium* samples were extracted with the Spectrum Total Plant RNA kit (Sigma) following the manufacturer protocol. In total, 17 *Arabidopsis*, 20 *Arachis*, 15 *Chenopodium*, and 20 *Gossypium* samples were extracted for RNAseq (Supplementary Table 1). RNA was quantified using the Agilent 2100 BioAnalyzer and sent to the Yale Center for Genome Analysis (YCGA) for library construction and sequencing. Illumina libraries were constructed using the TruSeq Stranded Total RNA kit with Ribo-Zero Plant and sequenced on a NovaSeq 6000 S4 flow cell. A minimum of 40 million read pairs (2 x 150 nt) was generated for each sample. Raw sequencing reads are available through the Short Read Archive (SRA) under PRJNA726938.

### Reference preparation and RNA-seq processing

Reference sequences for each genus were prepared by concatenating primary transcripts for each polyploid species with transcripts for each organelle (Supplementary Table 2). Primary transcripts were derived from recent genome sequences published for *Arabidopsis suecica* (Novikova et al. 2017), *Arachis hypogea* (Bertioli *et al*. 2019), *Chenopodium quinoa* (Jarvis *et al*. 2017), and *Gossypium hirsutum* (Chen *et al*. 2020). RepeatMasker (Smit *et al*. 2015) was used to mask each set of nuclear primary transcripts with both the organellar genomes and transcriptomes (Supplementary Table 2, and see below) for each species, and any transcript with fewer than 75 nucleotides of non-organelle derived sequence was discarded. Mitochondrial and plastid transcripts for each genus were derived from publicly available organelle genome annotations for a single representative species from each genus (Supplementary Table 2), with the exception of *Arachis* mitochondrial genes (see below). Each protein-coding gene set was manually curated to (1) add genes that were absent from the GenBank annotations (via BLAST identification; (Camacho *et al*. 2009)), (2) remove duplicate gene copies from the plastid inverted repeat, (3) remove non-conserved hypothetical genes, and (4) standardize gene naming conventions. Because there is no complete mitochondrial genome published for any *Arachis* species, we used available transcriptomic and genomic resources to extract protein-coding sequences for *Arachis* mitochondrial genes. Most genes were recovered by performing tBLASTN of *Arabidopsis* protein sequences against an unpublished dataset of *Arachis hypogaea* full-length cDNAs generated with PacBio Iso-Seq technology (NCBI Sequence Read Archive accession SRR14414925), and the remaining mitochondrial genes were extracted by searching against *Arachis hypogaea* genomic contigs in PeanutBase (Dash *et al*. 2016). Our curated mitochondrial and plastid protein-coding reference sequences for each taxon are available via https://github.com/Wendellab/CytonuclearExpression.

RNA-seq reads for each species were processed via Kallisto v0.46.1 (Bray *et al*. 2016) (i.e., *kallisto quant*) to assign orthologs and/or homoeologs to genes and quantify transcripts. Following Kallisto quantification, a principal component analysis (PCA) was generated for each genus using SNPRelate (Zheng *et al*. 2012) in R/4.0.2 (R Core Team 2020) to verify sample identity and generate an overview of the count data. PCA plots were visualized using ggplot2 (Wickham 2016) in R. Clustering heatmaps were generated using pheatmap (Kolde 2012). Code pertaining to this project can be accessed at https://github.com/Wendellab/CytonuclearExpression.

### Ortholog identification and targeting inference

We followed the methods of (Sharbrough *et al*. 2021) to identify orthologous genes arising from allopolyploidy (*i.e.*, ‘quartets’ consisting of one homolog from each diploid parent and two homoeologs from the allopolyploid). Briefly, we used Orthofinder (v2.3.8) (Emms and Kelly 2019) to cluster protein coding genes into homologous gene families. We retained orthogroups containing three or more homologs, extracted coding sequences (CDS) for those proteins, and aligned each using the L-INS-i algorithm in MAFFT (v7.480) (Katoh and Standley 2013). Model selection was done using jModelTest2 v2.1.10 (Darriba *et al*. 2012) and phylogenetic inference was performed in PhyML v3.3.20211021 (Guindon and Gascuel 2003), as previously described (Sharbrough *et al*. 2021). Because these gene trees often contain multiple orthologous groups resulting from ancient duplications, we extracted subtrees containing potential quartets (*i.e.* subtrees with the expected number of genes from each species) using *<u>subTreeIterator.py</u>* (Sharbrough *et al*. 2021). We merged these phylogenetically-based quartet predictions with independent synteny-based quartet predictions (generated via pSONIC; (Conover *et al*. 2021)) to identify high-confidence quartets. Quartets that were predicted by at least one method and were not in conflict with the second method were retained for analysis. Each quartet was analyzed for organelle targeting information using combined information from (1) CyMIRA (Forsythe *et al*. 2019); (2) *de novo* targeting software, including iPSORT v0.94 (Bannai *et al*. 2002), LOCALIZER v1.0.4 (Sperschneider *et al*. 2017), Predotar v1.03 (Small *et al*. 2004), and TargetP v1.1b (Emanuelsson *et al*. 2007); and (3) Orthofinder-based homology to the *Arabidopsis thaliana* Araport 11 proteome. Full details can be found in (Sharbrough *et al*. 2021), and relevant scripts can be found at https://github.com/jsharbrough/CyMIRA_gene_classification and https://github.com/jsharbrough/allopolyploidCytonuclearEvolutionaryRate/blob/master/scripts/su <u>bTreeIterator.py</u>, as well as https://github.com/Wendellab/CytonuclearExpression.

### Differential gene expression

Differential gene expression analyses were conducted in R/4.0.2 (R Core Team 2020) using DESeq2 (Love *et al*. 2014) with the design ‘∼species’ and with the reference transcriptomes detailed above. Genes with a Benjamini–Hochberg (Benjamini and Hochberg 1995) adjusted p-value <0.05 (as implemented in DESeq2) were considered differentially expressed. Expression PCA and pheatmaps were made in R using the base R package and pheatmap v1.0.12.

Differential expression (DE) was evaluated three ways: (1) DE between diploid progenitor and corresponding polyploid subgenome, (2) DE between each diploid progenitor and the total polyploid expression (*i.e.,* summed homoeolog expression), and (3) DE between maternal and paternal homoeologs. Enrichment of differential expression (DE) genes in cytonuclear gene categories was conducted using Fisher’s Exact Test (*fisher.test*) relative to the not-organelle-targeted (NOT) category.

We employed a mixed-effects modeling approach to test whether differences in expression across homoeologs were related to cytonuclear targeting category (inferred from CyMIRA), legacy effects of diploid progenitors (estimated here as the difference in expression across diploid relatives), and the interaction between targeting category and legacy effects. Expression modeling was conducted in R/4.1.1 and considered the two models: (1) *Δrlog* ∼ Targeting, and (2) *Δrlog* ∼ Targeting + ΔrlogDiploid + Targeting × *Δrlog*_Diploid_, where *Δrlog* represents the difference in DESeq2-derived rlog normalized counts (maternal - paternal homoeolog), ΔrlogDiploid represents the difference in DESeq2-derived rlog normalized counts between the model diploid progenitors, Targeting represents the CyMIRA identified targeting category, and Targeting × *Δrlog*_Diploid_ represents the interaction between category and diploid expression levels. Fixed effects for each model were evaluated using emmeans v1.7.0 and the analysis of variance (ANOVA) was evaluated using car v3.0-11, with a type II computation of the sums-of-squares. Because model 1 is nested within model 2, we compared these two models for each species using lrtest from lmtest v0.9-39 in R/4.1.1.

### Functional enrichment tests

CyMIRA-based results were verified for *Arabidopsis suecica, Gossypium hirsutum,* and *Gossypium barbadense* using FUNC-E in conjunction with existing functional annotations from INTERPRO (Jones *et al*. 2014), GO ontology (The Gene Ontology Consortium 2019), and Plant Ontology (available for *Arabidopsis* only; (Avraham *et al*. 2008)). *Arabidopsis* functional annotations were downloaded from TAIR (Cheng *et al*. 2017), and the *Gossypium* functional annotations were downloaded from CottonFGD (Zhu *et al*. 2017), both accessed in January 2022. These custom ontology lists were used to generate vocabulary terms for each FUNC-E analysis (one per species). Two sets of genes were used as queries in functional enrichment analyses, both of which are restricted to ortholog-homoeolog quartets with statistically significant differential expression between homoeologs (DESeq2 p-value < 0.05) that was also greater than fourfold. An additional criterion for the second query gene set required that the difference in fold change (FC) between homoeologs and FC between parental orthologs also had to be greater than four (*i.e.*, | ΔFC | > 4). In both cases, the reference (*i.e.*, background) set was composed of quartets regardless of p-value and/or fold-change; these comprised 11,307 for *Arabidopsis suecica*, 18,669 for *Gossypium hirsutum*, and 18,099 for *Gossypium barbadense*.

Functional enrichment was determined in FUNC-E via a one-sided Fisher’s Exact Test for each comparison, and multiple tests were subjected to Benjamini correction; significance was determined as adjusted p < 0.05. By default, upregulated and downregulated genes were tested separately.

## Results

### Generation and categorization of reference sequences

Representative transcriptomes for each genus were downloaded along with both organellar genomes and transcriptomes (Supplementary Table 2). In the case of *Arachis*, only putative transcripts were available for the mitochondria (see methods). Because reference genomes frequently have nuclear insertions of organellar genes that can be included in predicted transcripts, we first masked each nuclear transcriptome (primary transcripts only) with both the matching organellar genomes and transcriptomes, and we subsequently removed transcripts with fewer than 75 surviving nucleotides. Between 206 and 2,510 nuclear transcripts were filtered from each reference, leaving between 44,175 and 73,595 non-organellar nuclear transcripts.

These were combined with the curated organellar transcriptomes, consisting of 108 - 112 genes in total (see methods), resulting in polyploid reference transcriptomes ranging from 44,283 to 73,707 genes (Table 1).

**Table 1:** Composition of the mapping reference for each genus. Primary transcripts from each nuclear transcriptome were masked using the organellar transcriptomes and genomes, and nuclear transcripts matching organellar sequences were removed. Gene quartets composed of a single gene for each diploid species and two paired homoeologs from the polyploid reference were identified. Each quartet was classified with respect to putative organellar targeting. “Dual-targeted” transcripts are those that have targeting information for both organelles. “Interacting” transcripts code for products that interact with organellar gene products, whereas “non-interacting” transcripts are those which function in one or both organelles but do not physically interact with an organellar gene product.

|  | *Arabidopsis* | *Arachis* | *Chenopodium* | *Gossypium* |
| --- | --- | --- | --- | --- |
| **mitochondrial transcripts** | 32 | 32 | 30 | 35 |
| **chloroplast transcripts** | 78 | 76 | 78 | 77 |
| **nuclear transcripts** | 44,625 | 67,150 | 44,770 | 74,902 |
| nuclear transcripts, excluding norgs | 44,419 | 64,640 | 44,175 | 73,595 |
| **total transcripts** | 44,735 | 67,258 | 44,878 | 75,014 |
| removed genes | 206 | 2,510 | 595 | 1,307 |
| ***total transcripts, excluding norgs*** | 44,529 | 64,748 | 44,283 | 73,707 |
| **Homoeologous pairs (genome)** | 12,254 | 11,671 | 9,231 | 20,124 |
| Not Targeted | 9,830 | 10,121 | 7,575 | 17,606 |
| Dual-targeted, interacting | 45 | 52 | 62 | 76 |
| Dual-targeted, non-interacting | 185 | 746 | 771 | 1,103 |
| Mitochondria-targeted, interacting | 263 | 156 | 169 | 326 |
| Mitochondria-targeted, non-interacting | 467 | 94 | 84 | 135 |
| Plastid-targeted, interacting | 168 | 133 | 159 | 246 |
| Plastid-targeted, non-interacting | 1,296 | 369 | 411 | 632 |

A curated set of high-confidence homoeologs was generated for each reference genome using a combination of phylogenetics and synteny (see methods), which were subsequently characterized by their potential to interact with either/both organelles (Table 1). The number of homoeologous pairs in each genome ranged from 9,231 in *Chenopodium quinoa* to 20,124 in *Gossypium hirsutum*, representing twice that number of genes (18,462 and 40,248 homoeologs, respectively). As expected, most genes (80-87%) were not predicted to be targeted to either organelle, with an average of 2-3% of genes placed in the six organelle-related categories (*i.e.*, mitochondria-/plastid-/dual-targeted, interacting/non-interacting genes; range = 0 - 11%; Table 1), as determined by CyMIRA (see methods). Of those genes exhibiting signatures of organelle targeting, homoeolog pairs that function in the organelle but do not have direct interactions with organellar-encoded proteins were generally more abundant, with the exception of mitochondria-targeted interacting genes, which were 1.5 - 2 times more abundant in most species (except *Arabidopsis thaliana*; Table 1) than the non-interacting mitochondrial genes. These targeting predictions were subsequently applied to the reference transcriptome generated for each genus (Table 1 and see methods).

We also evaluated the degree of homoeolog loss between the maternal and paternal genome for genes where orthologs were recovered from both model progenitors but only one polyploid subgenome (Table 2). If there is a general cytonuclear incompatibility between the diploid progenitors, then we would expect an excess in paternal homoeolog loss for genes involved in cytonuclear categories, *i.e.*, dual-targeted interacting (DI), dual-targeted non-interacting (DNI), mitochondria-targeted interacting (MI), mitochondrial-targeted non-interacting (MNI), plastid-targeted interacting (PI), and plastid-targeted non-interacting (PNI). Because the *Chenopodium quinoa* has a large number of genes not assigned to maternal/paternal subgenome, and the *Arachis hypogea* genome exhibits a high degree of homoeologous exchange (thereby reducing the number of reliable quartets), we restricted our analysis of putative homoeolog loss to *Arabidopsis suecica* and *Gossypium hirsutum* (Table 2). For most categories, there was no significant difference in paternal versus maternal homoeolog loss relative to background (*i.e.,* genes whose products are not targeted to either organelle (NOT); Fisher’s Exact *p* > 0.05). Only one cytonuclear category from the two genomes (*i.e.,* DNI from *Arabidopsis*) exhibited biased homoeolog loss, and the distribution of loss was contrary to what is expected given maternal inheritance of organelles.

**Table 2.** The number of paternal or maternal homoeologs lost from *Arabidopsis* and *Gossypium* for each category, and the proportion that represent maternal losses. If broad cytonuclear incompatibilities exist, we expect that the number of maternal homoeologs lost should be fewer in cytonuclear gene categories than for the rest of the genome, represented by low numbers in the %maternal columns. Cytonuclear categories that are statistically different in distribution from Non-organelle-targeted (NOT) genes are marked with an * in the column where loss is greater than expected by the NOT category (Fisher’s exact p <0.05).

|  | *Arabidopsis* |  |  | *Gossypium* |  |  |
| --- | --- | --- | --- | --- | --- | --- |
|  | paternal loss | maternal loss | % maternal | paternal loss | maternal loss | % maternal |
| Not-organelle-targeted | 673 | 439 | 39% | 342 | 542 | 61% |
| All Cytonuclear | 106 | 81 | 46% | 21 | 30 | 59% |
| Dual-targeted_ Interacting | 2 | 3 | 60% | 0 | 2 | 100% |
| Dual-targeted_ Non-interacting | 2 | 8* | 80% | 9 | 13 | 59% |
| Mitochondria-targeted_ Interacting | 19 | 10 | 34% | 4 | 4 | 50% |
| Mitochondria-targeted_ Non-interacting | 20 | 16 | 44% | 1 | 2 | 67% |
| Plastid-targeted_ Interacting | 12 | 7 | 37% | 1 | 2 | 67% |
| Plastid-targeted_ Non-interacting | 51 | 45 | 47% | 6 | 7 | 54% |

### Sequencing yields and general gene expression

Because the aim of this study was to characterize cytonuclear accommodation at the level of gene expression in polyploid species, total RNA was extracted for each accession and ribodepletion was used to remove ribosomal RNAs, circumventing the bias of polyA-selection protocols that exclude some organellar transcripts (Slomovic *et al*. 2006, 2008; Smith 2013). As expected, transcripts from the organelles were abundant <u>Supplementary 3</u>; however, sufficient nuclear transcriptome coverage was achieved, ranging from 26 to 90 M reads per sample (averages are *Arabidopsis* = 61 M, *Arachis* = 36 M, *Chenopodium* = 44 M, *Gossypium* = 61 M). One replicate each for *Arachis hypogea* and *Arachis* IpaDur1 was removed due to low mapping rates (i.e., < 25% of reads mapped; averages without outliers are *Arabidopsis* = 79%, *Arachis* = 82%, *Chenopodium* = 62%, and *Gossypium* = 72% of reads mapped). PCA and hierarchical clustering of the gene expression data exhibit clustering of replicates for each species within a genus, with one exception. *Chenopodium suecicum* replicate #1 was placed intermediate among all *Chenopodium* species via PCA (Supplementary Figure 1), and it clustered with *Chenopodium quinoa* via hierarchical clustering. Because this sample may represent a contaminated hybrid, it was excluded from subsequent analyses.

In general, the polyploid species exhibited more up-regulated genes than down-regulated genes relative to their diploid counterparts, both with respect to homoeolog-progenitor comparisons and total polyploid expression (Table 3). This pattern was most prominent in cotton, where all comparisons exhibited more up-regulated than down-regulated genes in polyploids (chi^2^ p<0.05), followed by *Arabidopsis suecica*, where all maternal comparisons exhibited more up-regulation. Conversely, *Chenopodium quinoa* only exhibited more up-regulation of the total polyploid expression (*i.e.*, the summed expression of homoeologs), and the natural peanut polyploid, *Arachis hypogea*, only exhibited more up-regulation of maternal homoeologs relative to expression in the model maternal diploid progenitor, *Arachis duranensis* (Table 3). Interestingly, the synthetic allotetraploid, *Arachis* IpaDur1 also exhibits more up-regulation of *Arachis duranensis* homoeologs, here functioning as the paternal diploid progenitor, with concomitant down-regulation in expression of homoeologs from the maternal diploid parent, *Arachis ipaensis*, potentially indicating a general bias toward *Arachis duranensis* expression.

**Table 3.** The total number of genes passing filter (see methods), and the number that are differentially expressed (parsed as up or down regulated). Cells that are highlighted are significantly different from equal (up-regulation vs down-regulation); chi2 < 0.05). Note that the different number of genes in the diploid-polyploid comparison (parsed as homoeologs) reflect differences in survivability in the DE analysis.

|  |  | *Gossypium hirsutum* |  | *Gossypium barbadense* |  | *Chenopodium quinoa* |  | *Arabidopsis suecica* |  | *Arachis hypogea* |  | *Arachis IpaDur1* |  |
| --- | --- | --- | --- | --- | --- | --- | --- | --- | --- | --- | --- | --- | --- |
| diploid divergence |  | 5 - 10 mya |  |  |  | 11 mya |  | 6 - 8 mya |  | 2.2 mya |  |  |  |
| polyploid origin |  | 1 - 2 mya |  |  |  | 2.5 - 3 mya |  | 16 kya |  | 9,400 ya |  | synthetic |  |
|  |  | A | D | A | D | A | B | T | A | D | I | I | D |
| diploid-polyploid,<br>parsed as homoeologs | Total genes | 18,197 | 18,355 | 18,197 | 18,355 | 8,603 | 8,616 | 11,154 | 11,225 | 10,154 | 10,404 | 10,404 | 10,154 |
|  | down-regulated | 2,803 (15%) | 3,912 (21%) | 2,617 (14%) | 3,461 (19%) | 2,390 (28%) | 1,984 (23%) | 1,358 (12%) | 1,888 (17%) | 328 (3%) | 1,778 (17%) | 2,290 (22%) | 28 (0%) |
|  | up-regulated | 3,166 (17%) | 4,390 (24%) | 2,908 (16%) | 4,052 (22%) | 2,519 (29%) | 1,965 (23%) | 1,478 (13%) | 1,987 (18%) | 593 (6%) | 1,716 (16%) | 1,868 (18%) | 166 (2%) |
| diploid-polyploid, total<br>expression in<br>polyploids versus one<br>or the other diploid | Total genes | 18,792 | 18,792 | 18,792 | 18,792 | 8,813 | 8,813 | 11,610 | 11,610 | 10,645 | 10,645 | 10,645 | 10,645 |
|  | down-regulated | 3,387 (18%) | 4,012 (21%) | 3,133 (17%) | 3,529 (19%) | 2,322 (26%) | 2,435 (27%) | 1,828 (16%) | 2,040 (18%) | 669 (6%) | 1,722 (16%) | 1,617 (15%) | 90 (1%) |
|  | up-regulated | 4,085 (22%) | 4,691 (25%) | 3,944 (21%) | 4,351 (23%) | 2,551 (29%) | 2,398 (27%) | 2,160 (19%) | 2,086 (18%) | 609 (6%) | 1,862 (18%) | 1,893 (18%) | 108 (1%) |

### Expression level dominance in cytonuclear genes

Expression level dominance (ELD) is a phenomenon whereby the combined expression of homoeologs in a polyploid is statistically similar to one diploid parent and statistically dissimilar from the other parent. In the context of cytonuclear compatibility, we might expect a bias toward the maternal diploid expression level (*i.e.*, ELD) for the combined expression of both homoeologs in cytonuclear gene categories. When we consider expression level dominance of nuclear genes within each species, irrespective of category (*i.e.,* NOT or any cytonuclear category), we see a general bias towards maternal ELD for *Arachis hypogea* and both species of *Gossypium* (binomial test, *p* < 0.05), but not for *Arabidopsis suecica* or *Chenopodium quinoa* (binomial test, *p* > 0.05; Table 4; Supplementary Table 4). These results are also reflected in the NOT category itself, where *Arachis hypogea* and both *Gossypium* species exhibit bias toward maternal ELD. Interestingly, however, when we compare patterns of ELD for all organelle targeted genes versus those in the NOT category, we find that *Arachis hypogea* and *Chenopodium quinoa* have significantly more genes (Fisher’s exact, p<0.05) exhibiting ELD in maternally-biased categories (*i.e.,* categories IV and IX; Supplementary Table 4) than expected from the overall distribution of maternal and paternal ELD, whereas both species of *Gossypium* exhibited similar patterns of ELD for cytonuclear genes as NOT genes. Notably, *Arachis* IpaDur1 exhibited an excess of paternal ELD, which is in contrast to the maternal ELD exhibited by *Arachis hypogea* but biased toward the same diploid parent (*i.e.,* biased toward *Arachis duranensis* in both cases). On the level of individual categories, four categories in three species exhibit an excess of ELD (Fisher’s exact, *p* < 0.05), all maternally biased: *Arachis hypogea*, DNI; *Chenopodium quinoa*, DNI and MI; and *Gossypium barbadense*, PI. All other individual categories exhibited similar ELD bias as displayed by the NOT genes for that species (Table 4).

**Table 4.** Number of genes exhibiting expression level dominance (ELD) toward each parental expression level, parsed by cytonuclear category. Categories are dual-targeted interacting (DI), dual-targeted non-interacting (DNI), mitochondria-targeted interacting (MI), mitochondrial-targeted non-interacting (MNI), plastid-targeted interacting (PI), and plastid-targeted non-interacting (PNI).

|  | *Arabidopsis suecica* |  | *Arachis hypogea* |  | *Arachis hypogea* |  | *Chenopodium quinoa* |  | *Gossypium barbadense* |  | *Gossypium hirsutum* |  |
| --- | --- | --- | --- | --- | --- | --- | --- | --- | --- | --- | --- | --- |
| Category | Maternal ELD | Paternal ELD | Maternal ELD† | Paternal ELD | Maternal ELD | Paternal ELD† | Maternal ELD | Paternal ELD | Maternal ELD† | Paternal ELD | Maternal ELD† | Paternal ELD |
| Not-organelle-targeted | 1481 | 1486 | 1659 | 365 | 41 | 2026 | 1294 | 1293 | 2961 | 2421 | 3063 | 2273 |
| All cytonuclear | 419 | 409 | 361* | 54 | 1 | 420 | 361* | 288 | 430 | 345 | 462 | 325 |
| DI | 8 | 8 | 11 | 1 | 0 | 11 | 15 | 14 | 9 | 12 | 12 | 15 |
| DNI | 31 | 30 | 184* | 25 | 0 | 203 | 184* | 126 | 194 | 156 | 213 | 156 |
| MI | 51 | 39 | 23 | 8 | 0 | 39 | 42* | 21 | 47 | 46 | 54 | 33 |
| MNI | 85 | 82 | 26 | 4 | 1 | 28 | 10 | 14 | 23 | 26 | 29 | 19 |
| PI | 22 | 37 | 32 | 2 | 0 | 38 | 31 | 29 | 48* | 21 | 46 | 30 |
| PNI | 222 | 213 | 85 | 14 | 0 | 101 | 79 | 84 | 109 | 84 | 108 | 72 |
| † parental ELD is significantly different from 1:1 and biased toward the noted parent |  |  |  |  |  |  |  |  |  |  |  |  |
| * cytonuclear category distribution (maternal versus paternal) is significantly different from the distribution in the NOT category and overrepresented by the noted parent |  |  |  |  |  |  |  |  |  |  |  |  |

We also identified some genes in these polyploids with expression levels that fell outside the range of the two parental diploid models (*i.e.*, transgressive expression), which may be associated with organelle copy number in a cytonuclear context. When considering all genes, regardless of targeting, *Arabidopsis suecica* and both *Arachis* species have statistically similar numbers of genes that are transgressive down-regulated (categories III, VII, and X in Supplementary Table 4) as transgressive up-regulated (categories V, VI, and VIII), whereas *Chenopodium quinoa* and both species of *Gossypium* have ∼20-35% more genes exhibiting transgressive up-regulation (versus down-regulation; Supplementary Table 4). Accounting for these global patterns, we find no species-category combinations exhibiting transgressive expression patterns in cytonuclear genes that are statistically different from NOT genes (after Benjamini-Hochberg *p*-value correction for multiple testing), although we note that many of these cytonuclear categories had very few genes (Supplementary Table 4) and are therefore difficult to statistically characterize.

### Homoeolog expression in cytonuclear genes

We evaluated homoeolog expression for each polyploid species in the context of the six cytonuclear categories with the biological expectation that maternal homoeologs should be preferentially up-regulated relative to paternal homoeologs (Figure 1 and Supplementary Tables 5-8). Figure 1 summarizes the results of the homoeolog comparisons for each homoeolog in 2 x 2 grids for each species-category, where maternal (left) and paternal (right) expression is measured relative to the model diploid progenitor and over-/under-representation is determined relative to the pattern observed in NOT genes (*i.e.*, background). Because cytonuclear incompatibility predicts upregulation of the co-evolved maternal cytonuclear homoeologs and down-regulation of the evolutionarily more distant paternal homoeologs, we expect a combination of the following patterns (Figure 1): (1) overrepresentation (depicted in red) for maternal homoeolog up-regulation (upper left square), (2) overrepresentation (red) for paternal homoeolog down-regulation (lower right square), (3) underrepresentation (depicted in blue) for maternal homoeolog down-regulation (lower left square), and/or (4) underrepresentation (blue) for paternal homoeolog up-regulation (upper right square). In general, fewer than half of the categories per polyploid species are consistent with cytonuclear incompatibility expectations, and, in both *Arachis* IpaDur1 and *Gossypium barbadense*, we do not observe any categories whose patterns are consistent with our biological expectations. None of the categories were consistent with cytonuclear expectations in more than two species, although each category was significant in at least one. Interestingly, the most frequently observed patterns were contrary to cytonuclear expectations (Figure 1); that is, 12 species-category comparisons contradict cytonuclear expectations (versus 7 consistent species-categories), although these contradictory patterns were also observed in no more than half of the categories per species.

**Figure 1.**
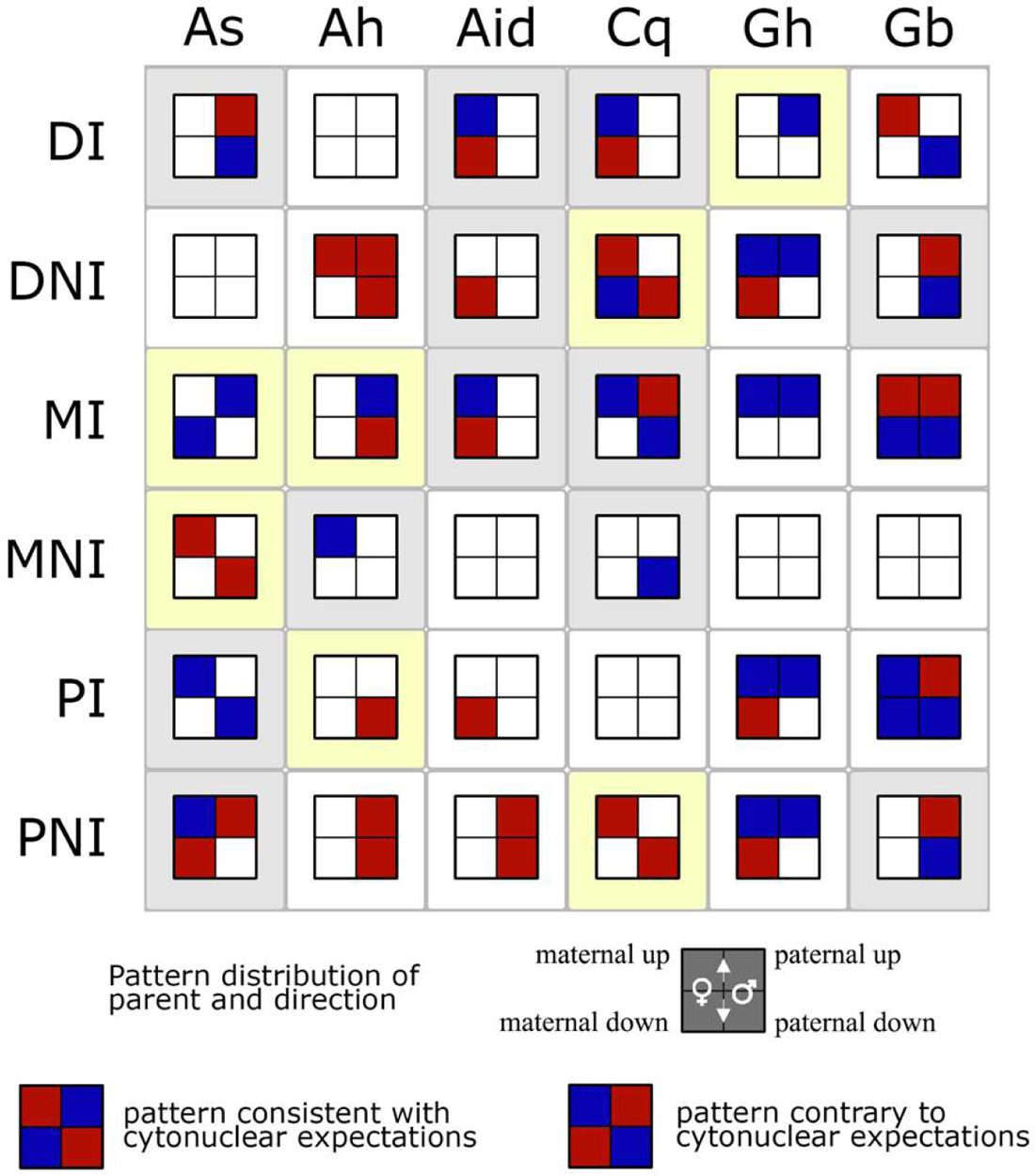
Summary of differential gene expression in cytonuclear categories for each polyploid species relative to each model diploid progenitor, partitioned as homoeologs. This pictogram displays the statistically significant (Fisher’s exact *p* < 0.05) overrepresentation (red) or underrepresentation (blue) of up- or down-regulated genes for each category, relative to non-cytonuclear genes. Each species/category is represented by a four-square grid, where the rows specify regulation (up or down) and columns specify the homoeolog comparison (i.e., maternal homoeolog vs maternal progenitor and paternal homoeolog versus paternal progenitor, respectively). In each quadrant, red indicates that there were more genes statistically significant in that parent-category combination than was expected based on the NOT distribution, whereas blue indicates there were fewer statistically significant genes in that parent-category combination. Example color patterns consistent with and contrary to cytonuclear expectations are shown on the bottom. Species-category combinations highlighted in yellow are consistent with the hypothesis that cytonuclear accommodation in polyploid species favors expression from the “more compatible” maternal genome (via up-regulation) and/or diminishes expression from the potentially “less favorable” paternal genome (via down-regulation), whereas species-category highlighted in grey specifically contradict cytonuclear expectations. Species include *Arabidopsis suecica* (As), *Arachis hypogea* (Ah), *Arachis* IpaDur1 (Aid), *Chenopodium quinoa* (Cq), *Gossypium hirsutum* (Gh), and *Gossypium barbadense* (Ah). Categories include Dual-Targeted Interacting (DI), Dual-Targeted Non-Interacting (DNI), Mitochondria-Targeted Interacting (MI), Mitochondria-Targeted Non-Interacting (MNI), Plastid-Targeted Interacting (PI), Plastid-Targeted Non-Interacting (PNI). All comparisons are relative to the Non-Organelle Targeted (NOT) genes.

**Table 5.** Homoeolog expression biases for each polyploid, partitioned as maternal and paternal bias. Bias is considered when homoeolog expression is statistically significant (adjusted p< 0.05), regardless of the magnitude of the change. The distribution of maternally-paternally biased genes for each cytonuclear category was evaluated relative to the NOT category using a Fisher’s Exact Test. Significant deviations (p<0.05) from the NOT distribution are noted by an asterisk, and the column (maternal or paternal) designates the parental bias that is overrepresented for that category.

|  | *Arabidopsis suecica* |  | *Arachis hypogea* |  | *Arachis IpaDur* |  | *Chenopodium quinoa* |  | *Gossypium hirsutum* |  | *Gossypium barbadense* |  |
| --- | --- | --- | --- | --- | --- | --- | --- | --- | --- | --- | --- | --- |
|  | Maternal bias | Paternal bias | Maternal bias | Paternal bias | Maternal bias | Paternal bias | Maternal bias | Paternal bias | Maternal bias | Paternal bias | Maternal bias | Paternal bias |
| Total | 1634 | 1887† | 757 | 836† | 2282† | 1678 | 1376 | 1613† | 3282 | 3690† | 2536 | 2734† |
| NOT | 1251 | 1527 | 628 | 689 | 1878 | 1397 | 1088 | 1310 | 2836 | 3151 | 2262 | 2345 |
| DI | 5 | 4 | 3 | 3 | 11 | 12 | 10 | 9 | 22* | 13 | 7 | 8 |
| DNI | 33 | 29 | 64 | 74 | 198 | 136 | 125 | 142 | 184 | 231 | 127 | 172* |
| MI | 53 | 53 | 6 | 11 | 36 | 24 | 16 | 35* | 64 | 73 | 24 | 36 |
| MNI | 67 | 75 | 8 | 9 | 24 | 20 | 18 | 14 | 23 | 29 | 14 | 24 |
| PI | 28* | 17 | 13 | 18 | 46 | 26 | 37* | 22 | 41 | 55 | 27 | 47* |
| PNI | 197* | 182 | 35 | 32 | 89 | 63 | 82 | 81 | 112 | 138 | 75 | 102* |
| † parental HEB is significantly different from 1:1 and biased toward the noted parent |  |  |  |  |  |  |  |  |  |  |  |  |
| * cytonuclear category distribution (maternal versus paternal) is significantly different from the distribution in the NOT category and overrepresented by the noted parent |  |  |  |  |  |  |  |  |  |  |  |  |

We also directly compared expression between homoeologs to ascertain the extent (or lack) of maternal expression bias, both in general and with respect to cytonuclear categories (Figure 2). Homoeolog expression bias (HEB) is distinct from expression level dominance (ELD) in that HEB reports statistically different expression levels *between* homoeologs, whereas ELD (see above) refers to instances where the *total* gene expression (of both homoeologs) is similar to only one parent. We find that most of the polyploids (except the synthetic *Arachis* IpaDur1) exhibit more genes with paternal HEB versus maternal, for all paired homoeologs regardless of category (Table 5). When these genes are partitioned into cytonuclear categories, however, we detect maternal bias for some individual categories, most notably *Arabidopsis suecica* and *Chenopodium quinoa*, where four of the six cytonuclear categories have more genes with maternal bias than paternal. In most cases, this directional shift toward maternal bias is not statistically significant from the NOT distribution (Fisher’s Exact Test, *p* > 0.05) and may either represent a lack of biological relevance or a lack of statistical power due to the small numbers in many of these categories (Table 5). The only categories that did exhibit statistically significant higher numbers of maternally HEB were the PI and PNI categories from *Arabidopsis suecica* and DI from *Gossypium hirsutum*. The latter may be somewhat surprising not only because this is the sole maternally biased category from either *Gossypium* species, but also because the closely related species *Gossypium barbadense* exhibits three cytonuclear categories with bias in the opposite direction (more paternal HEB than is expected from the NOT distribution, *i.e.,* DNI, PI, and PNI; Table 5).

**Figure 2.**
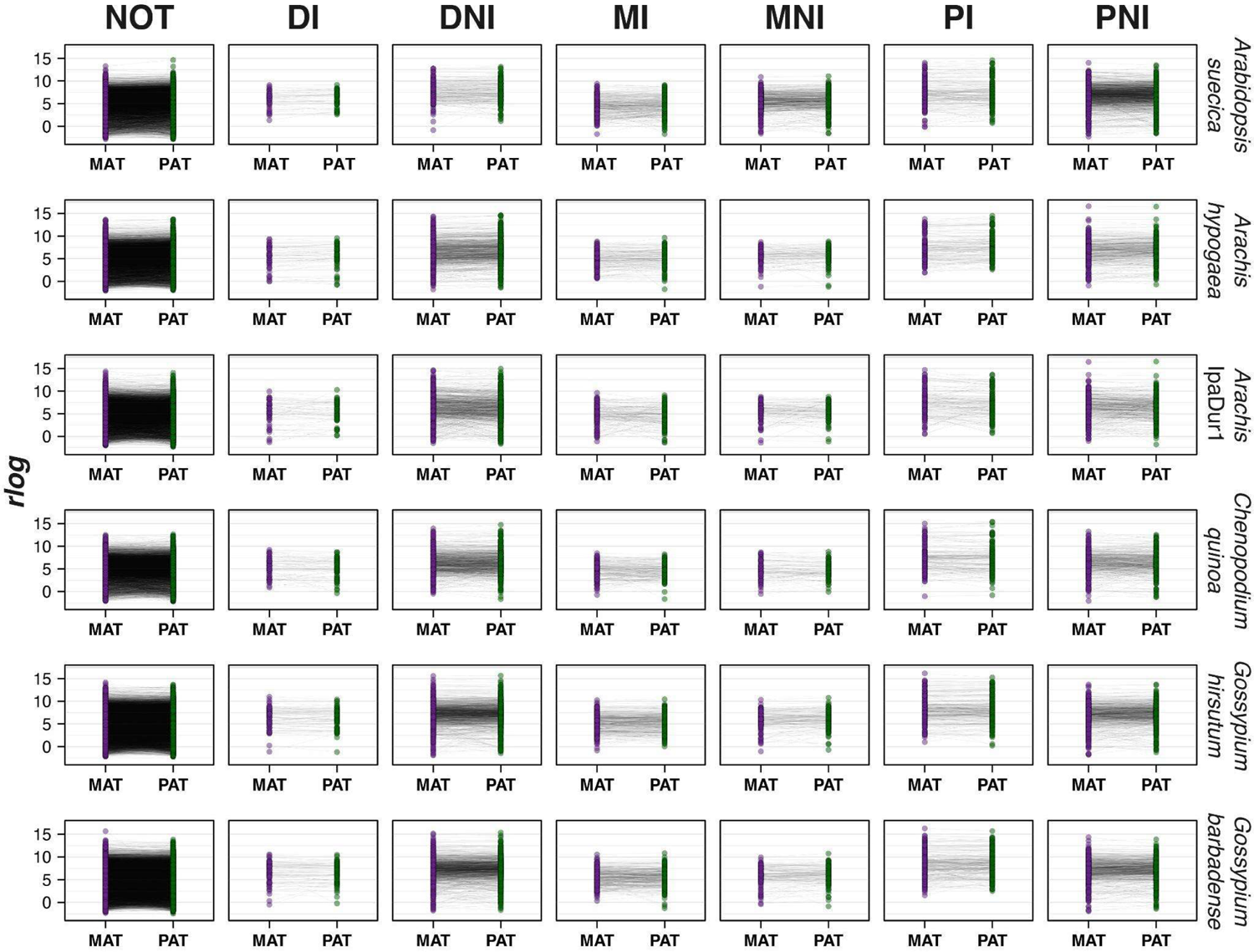
Mean normalized gene expression across homoeologs of six allotetraploids. Mean rlog values (circles) from 4-5 biological replicates each are depicted for maternal (left, purple) and paternal (right, green) homoeologs, partitioned into seven functional categories: Non-organelle-targeted (NOT), Dual-targeted Non-Interacting (DNI), Mitochondria-targeted Non-Interacting (MNI), Plastid-targeted Non-Interacting (PNI), Dual-targeted Interacting (DI), Mitochondria-targeted Interacting (MI), and Plastid-targeted Interacting (PI). Semi-transparent lines connect maternal and paternal homoeologs.

We further evaluated the possible effects of cytonuclear category membership on homoeolog expression using linear modeling. We began with a model that asked if the difference in observed expression between maternal and paternal homoeologs was a function of where it was targeted (*Δrlog* ∼ Targeting) using the six aforementioned categories. For this model, we evaluated expression in each polyploid as a difference in *rlog* normalized counts (derived from DESeq2) between the maternal homoeolog and the paternal homoeolog (as *Δrlog* = *rlog_Maternal_* - *rlog_Paternal_*). The results of this model (Table 6) suggest that membership in a cytonuclear category (*i.e.,* Targeting) does have an effect on the difference between homoeolog expression levels for *Arabidopsis suecica*, *Arachis* IpaDur1, *Chenopodium quinoa*, *Gossypium hirsutum,* and *Gossypium barbadense*, but it is not significant for *Arachis hypogea* (ANOVA, p <0.05).

**Table 6.** Type III ANOVA-based p-value for the *Targeting* category. Estimated Marginal Means effect size for individual contrasts between *Targeting* categories are listed below, with significant categories (p<0.05) marked with an *.

|  | *Arabidopsis suecica* | *Arachis hypogea* | *Arachis ipadur* | *Chenopodium quinoa* | *Gossypium hirsutum* | *Gossypium barbadense* |
| --- | --- | --- | --- | --- | --- | --- |
| Targeting (ANOVA p-value) | 1.13E-26* | 0.061 | 0.021* | 0.007* | 0.031* | 1.13e-05* |
| *Fixed Effects* |  |  |  |  |  |  |
| DI | 0.1006 | 0.1147 | -0.2537 | 0.1750 | 0.1351 | 0.1086 |
| DNI | 0.2202* | -0.0425 | -0.0151 | -0.0017 | -0.0041 | -0.0854* |
| MI | 0.1190* | -0.0713 | 0.0537 | -0.0500 | -0.0453 | -0.0451 |
| MNI | 0.0996* | 0.1271 | -0.3053* | 0.0756 | -0.1532* | -0.1351* |
| PI | 0.2209* | -0.1035 | 0.1383 | 0.1862* | -0.0459 | -0.1316* |
| PNI | 0.1871* | -0.0318 | -0.0401 | 0.0566 | -0.0237 | -0.0335 |
| *Contrasts* |  |  |  |  |  |  |
| NOT vs DI | -0.1006 | -0.1147 | 0.2537 | -0.1750 | -0.1351 | -0.1086 |
| NOT vs DNI | -0.2202* | 0.0425 | 0.0151 | 0.0017 | 0.0041 | 0.0854* |
| NOT vs MI | -0.1190 | 0.0713 | -0.0537 | 0.0500 | 0.0453 | 0.0451 |
| NOT vs MNI | -0.0996* | -0.1271 | 0.3053 | -0.0756 | 0.1532 | 0.1351 |
| NOT vs PI | -0.2209* | 0.1035 | -0.1383 | -0.1862* | 0.0459 | 0.1316 |
| NOT vs PNI | -0.1871* | 0.0318 | 0.0401 | -0.0566 | 0.0237 | 0.0335 |
| DI vs DNI | -0.1196 | 0.1572 | -0.2386 | 0.1767 | 0.1392 | 0.1940 |
| MI vs MNI | 0.0194 | -0.1983 | 0.3590 | -0.1256 | 0.1079 | 0.0900 |
| PI vs PNI | 0.0338 | -0.0717 | 0.1784 | 0.1297 | -0.0222 | -0.0982 |
| DI vs MI | -0.0184 | 0.1860 | -0.3074 | 0.2249 | 0.1804 | 0.1537 |
| DI vs PI | -0.1203 | 0.2182 | -0.3920 | -0.0113 | 0.1810 | 0.2403 |
| MI vs PI | -0.1019 | 0.0322 | -0.0846 | -0.2362 | 0.0006 | 0.0865 |
| DNI vs MNI | 0.1206 | -0.1696 | 0.2902 | -0.0773 | 0.1492 | 0.0497 |
| DNI vs PNI | 0.0331 | -0.0107 | 0.0250 | -0.0583 | 0.0196 | -0.0519 |
| MNI vs PNI | -0.0875 | 0.1589 | -0.2652 | 0.0190 | -0.1296 | -0.1017 |

The number and identities of categories with fixed effects significantly different from NOT vary between polyploids (Table 6; Supplementary Figure 2), with the MNI category exhibiting significant fixed effects most frequently (3 of 5 significant polyploids) while MI is not significant for any polyploid. Contrasts among categories are even less suggestive of expression differences due to targeting for most species, although in *Arabidopsis suecica* most categories (except DI and MI) exhibited significantly greater expression differences between homoeologs than the NOT category (*p* < 0.05) and in the expected direction (*i.e.,* expression differences between homoeologs in those cytonuclear categories are more maternally biased than NOT). In the remaining species, only PI in *Chenopodium quinoa* and DNI in *Gossypium barbadense* were significantly different from the NOT category, the latter of which contradicted our expectations (*i.e.,* NOT in *Gossypium barbadense* is more maternally biased than is DNI; Table 6).

Importantly, this first model fails to account for the effects of parental legacy on expression levels in the polyploid and how deviations from parental expression levels may occur within the polyploid, the latter of which may be important depending on functional category (Supplementary Figure 3). Therefore, we repeated the analysis with a second model that also considered the difference in diploid expression as an explanatory term for the observed difference in homoeolog expression (*i.e.*, *Δrlog* ∼ Targeting + *Δrlog_Diploid_* + Targeting × *Δrlog_Diploid_*). We find that both targeting category and legacy expression differences (*Δrlog_Diploid_*, representing the difference in the *rlog* values for the maternal and paternal diploid model species) both affect homoeolog expression differences and strongly interact (Targeting × *Δrlog_Diploid_*) in all comparisons (ANOVA, *p* < 0.05; Table 7). Unlike the previous model, all of the targeting categories are significant predictors of *Δrlog* in at least two polyploid species (Table 7). Additionally, contrasts in all species (except *Arachis hypogea*) suggest that two to four targeting categories per species are significant predictors of differences in homoeolog expression beyond that predicted by non-organelle-targeted genes (Table 7). Interestingly, however, the direction of these differences is not consistent and in some cases are contrary to the biological expectation that homoeolog expression differences will be more maternally biased in categories that interact with the maternally-inherited organelles. Here we find few instances of greater expression divergence between homoeologs in targeting categories (versus NOT), which are limited to most categories for *Arabidopsis suecica* and the DNI category in *Chenopodium quinoa* (Table 7). Conversely, three categories each in *Arachis* IpaDur1, *Gossypium hirsutum*, and *Gossypium barbadense* and one in *Chenopodium quinoa* (MI) exhibit a greater difference between homoeologs for the NOT category, which contradicts the assumption that organelle-targeted homoeologs should preferentially up-regulate maternal homoeologs and/or down-regulate paternal homoeologs, both of which increase the difference in expression between homoeologs.

**Table 7.** Type III ANOVA-based p-values for the categories *Targeting* and *dipDelta,* and their interaction term. Estimated Marginal Means effect size for individual contrasts between *Targeting* categories are listed below, with significant categories (p<0.05) marked with an *.

|  | *Arabidopsis suecica* | *Arachis hypogea* | *Arachis ipadur* | *Chenopodium quinoa* | *Gossypium hirsutum* | *Gossypium barbadense* |
| --- | --- | --- | --- | --- | --- | --- |
| Targeting | 6.35E-31* | 1.45E-04* | 1.53E-10* | 1.52E-10* | 1.63E-08* | 3.34E-14* |
| dipDelta | 0* | 0* | 0* | 0* | 0* | 0* |
| Targeting:dipDelta | 3.58E-17* | 7.17E-29* | 1.86E-32* | 4.89E-14* | 7.97E-05* | 6.98E-07* |
| *Fixed Effects* |  |  |  |  |  |  |
| DI | 0.2006 | 0.0326 | -0.4834* | -0.3167* | 0.1080 | 0.1288 |
| DNI | 0.3069* | -0.0800* | -0.1315* | 0.0344 | -0.0643* | -0.1225* |
| MI | 0.6809* | 0.0952 | -0.3517* | -0.3461* | 0.0674 | 0.0857* |
| MNI | 0.2744* | 0.1327 | -0.1239 | -0.1988* | -0.0095 | 0.0541 |
| PI | 0.4127* | -0.1174 | -0.1389 | 0.1174* | -0.1570* | -0.1893* |
| PNI | 0.1076* | -0.1208* | -0.0847 | -0.0398 | -0.0996* | -0.0883* |
| dipDelta | 0.3809* | 0.1800* | 0.6092* | 0.3879* | 0.4879* | 0.5021* |
| DI:dipDelta | 0.0556 | -0.1028* | -0.0122 | 0.1130* | -0.0330 | 0.0290 |
| DNI:dipDelta | 0.0765* | -0.0656* | 0.1965* | 0.0665* | -0.0226 | 0.0136 |
| MI:dipDelta | 0.1815* | 0.0675* | 0.3203* | 0.0686* | 0.1072* | 0.1285* |
| MNI:dipDelta | 0.0530* | 0.0482 | 0.2568* | 0.0626 | 0.0891* | 0.1377* |
| PI:dipDelta | 0.1723* | -0.0764* | 0.1347* | 0.0345 | -0.0486 | 0.0445 |
| PNI:dipDelta | 0.0066 | -0.0984* | 0.0136 | 0.1420* | -0.0023 | 0.0356* |
| *Contrasts* |  |  |  |  |  |  |
| NOT vs DI | -0.0742 | -0.1483 | 0.4824* | 0.2485 | -0.1329 | -0.1068 |
| NOT vs DNI | -0.1330* | 0.0062 | 0.1480* | -0.0748* | 0.0472 | 0.1327* |
| NOT vs MI | -0.2681* | -0.0192 | 0.3785* | 0.3047* | 0.0136 | 0.0113 |
| NOT vs MNI | -0.1538* | -0.0784 | 0.1454 | 0.1610 | 0.0768 | 0.0500 |
| NOT vs PI | -0.0209 | 0.0315 | 0.1502 | -0.1383 | 0.1202* | 0.2229* |
| NOT vs PNI | -0.0926* | 0.0101 | 0.0859 | -0.0464 | 0.0978* | 0.1152* |
| DI vs DNI | -0.0588 | 0.1545 | -0.3344 | -0.3233* | 0.1801 | 0.2396 |
| MI vs MNI | 0.1142 | -0.0592 | -0.2331 | -0.1437 | 0.0632 | 0.0386 |
| PI vs PNI | -0.0717 | -0.0214 | -0.0644 | 0.0919 | -0.0224 | -0.1078 |
| DI vs MI | -0.1939 | 0.1290 | -0.1038 | 0.0562 | 0.1465 | 0.1182 |
| DI vs PI | 0.0533 | 0.1797 | -0.3322 | -0.3868* | 0.2531* | 0.3298* |
| MI vs PI | 0.2472* | 0.0507 | -0.2283 | -0.4430* | 0.1067 | 0.2116* |
| DNI vs MNI | -0.0208 | -0.0846 | -0.0026 | 0.2358* | 0.0296 | -0.0828 |
| DNI vs PNI | 0.0404 | 0.0039 | -0.0621 | 0.0284 | 0.0507 | -0.0176 |
| MNI vs PNI | 0.0612 | 0.0885 | -0.0596 | -0.2074 | 0.0210 | 0.0652 |

### Tests of functional enrichment

Functional enrichment analyses were conducted for *Arabidopsis* and *Gossypium* to further assess whether the lack of clear cytonuclear patterns were also observable through broad functional categories (versus the heretofore used CyMIRA categorizations) for those species where suitable information was available. Using a list of species-specific vocabulary terms from existing resources (i.e., TAIR (Cheng *et al*. 2017) and CottonFGD (Zhu *et al*. 2017)) to annotate our gene sets, we compared the suite of genes with greater than four-fold differences in homoeolog expression (maternal *vs.* paternal) with those that exhibited any difference in homoeolog expression (regardless of fold-change or significance). More functional annotations are available in the model genus *Arabidopsis*, so it is unsurprisingly that a greater number of terms were enriched for *Arabidopsis* (126 terms; Supplementary Table 9) compared to *Gossypium* (75 and 52 terms for *Gossypium hirsutum* and *Gossypium barbadense*, respectively; Supplementary Table 10). Enriched terms in both *Arabidopsis suecica* and *Gossypium barbadense* were nearly evenly split with respect to parental bias, contrary to the general bias toward paternal homoeolog expression. In *Arabidopsis suecica*, 65 (out 126) terms exhibited paternal expression bias; likewise, 24 (out of 52) enriched terms exhibited paternal bias in *Gossypium barbadense*.

Conversely, *Gossypium hirsutum* exhibited a clear maternal bias in enriched terms, *i.e.,* 53 maternally-biased terms versus 22 paternally-biased (Supplementary Table 10). Of those terms exhibiting enrichment in DE genes (>fourfold change, relative to background), only *G. barbadense* contained organelle relevant terms (*i.e.*, GO:0009523, photosystem II; GO:0009654, photosystem II oxygen evolving complex; IPR002683, PsbP C-terminal; and GO:0015979, photosynthesis) all of which exhibited a general bias towards maternal expression (Supplementary Table 10). Because a given gene can have multiple Gene Ontology (GO) and/or InterPro (IPR) terms associated with it, these four vocabulary terms represent only 5 genes in *G. barbadense* with an average 4.9-fold difference between homoeologs. Notably, all four organelle relevant terms exhibited a general bias towards maternal expression (Supplementary Table 10), consistent with the biological expectation of preference for maternal cytonuclear genes.

Interestingly, although *Gossypium barbadense* had the only organelle-relevant terms, none of these remain enriched when the analysis is restricted to only those genes exhibiting more than a fourfold difference in expression between homoeologs *and* whose fold-change between homoeologs compared to fold-change between diploids was at least 4.0 (*i.e., ΔFC_Homoeolog_* > 4 & *ΔFC_Diploid_* > 4; see methods; Supplementary Table 10), possibly indicating that some of the observed differences are best explained by the diploid progenitors. Conversely, while no organelle related terms were enriched in *Arabidopsis suecica* when only homoeolog fold change (*ΔFC*_Homoeolog_ >4) was thresholded, a different functional term (*i.e.*, GO:0009941; “chloroplast envelope”) did show enrichment in the restricted set (*ΔFC_Homoeolog_* > 4 & *ΔFC_Diploid_* > 4).

Chloroplast envelope is associated with six pairs of maternally-biased, DE homoeologs whose average 10.6-fold difference in expression is substantially different from the average 0.6-fold difference in expression between parental orthologs. Interestingly, while chloroplast envelope alone is enriched here (and maternally-biased) for *Arabidopsis suecica*, the expression patterns in plastid-related CyMIRA categories (relative to the diploid parents) generally contrast our expectation of maternal up-regulation and/or paternal down-regulation (Figure 1).

### Expression accommodation in Rubisco

Previous analyses of the Rubisco small subunit (*rbcS*) cytonuclear gene family in multiple polyploid species reported patterns of maternally-biased gene conversion and preferential expression of maternally-converted paternal homoeologs (Gong *et al*. 2012, 2014); therefore, we specifically extracted expression patterns for rbcS from the current data. Consistent with previous results (Gong *et al*. 2012, 2014), we found that *rbcS* is composed of a small gene family in each polyploid species (<u>Table Rubisco</u>). Because our analyses are based on available genomic and/or transcriptomic reference sequences, which are far less developed for *Arachis hypogea*, we were unable to assign subgenomes (nor assess expression) for the six *rbcS* copies detected in either *Arachis* polyploid. For the remaining polyploids, the number of copies assigned to subgenome and/or paired as homoeologs varied depending on available information. In most cases where strict homoeologs could not be identified, it was due to copy number variation in the annotation. For example, the *Arabidopsis suecica* genome is divided into *Arachis thaliana* (maternal) and *Arabidopsis suecica* (paternal) contigs; however, seven of the nine *rbcS* copies are in two tandem arrays (AsAa_g20535-37 and AsAt_g19714-17), making orthology difficult to determine; the unpaired copies of *rbcS* in *Chenopodium* and *Gossypium* were also a result of tandem duplications complicating orthology assignment. In general, comparisons of *rbcS* between subgenome and diploid progenitor suggest upregulation of *rbcS* in the *Arabidopsis* and *Chenopodium*, but not in either *Gossypium* species (Table 8). Notably, the *Chenopodium rbcS* genes assigned to subgenome (*i.e.,* paternal: AUR62042566 and maternal: AUR62018154) follow our biological expectations in that the maternal homoeolog exhibits upregulation relative to the diploid state; however, a comparison of expression between these homoeologs suggests that the paternal homoeolog is expressed 1.4-fold greater than the maternal homoeolog, contrary to the expectation that the maternal homoeolog would be preferentially expressed. We also note that seven copies of *rbcS* were omitted from the *Chenopodium* analysis because they were not assignable to a subgenome, which may contribute to an overall bias that cannot be determined here. *Gossypium hirsutum*, on the other hand, exhibits a slight, but statistically significant, maternal homoeolog bias (Table 8), congruent with biological expectations; however, a similar limitation resulting in the omission of four *rbcS* copies may also affect our inferences in the present analysis.

**Table 8.** For *Arachis* and *Chenopodium*, incomplete information prohibited assignment of individual rbcS copies to subgenome (*i.e.,* n.d., or not determined). The average fold-change between each polyploid subgenome and it’s model diploid progenitor (e.g., Maternal/Paternal FC) is listed for each other species. Comparisons that did not achieve statistical significance are marked as NS. Differences in homoeolog expression for the single pair in each genome is listed (Homoeolog FC) and reported as paternal versus maternal.

|  | *A. suecica* | *A. hypogea* | *A. ipadur* | *C. quinoa* | *G. hirsutum* | *G. barbadense* |
| --- | --- | --- | --- | --- | --- | --- |
| **Copies** | 9 | 6 |  | 9 | 6 |  |
| **maternal** | 5 | n.d. |  | 1+ | 3 |  |
| **paternal** | 4 | n.d. |  | 1+ | 3 |  |
| **paired** | 1 (2 homoeologs) | 0 |  | 1 (2 homoeologs) | 1 (2 homoeologs) |  |
| Maternal FC | 2.7 | n.d. | n.d. | 5.2 | NS | NS |
| Paternal FC | 3.3 | n.d. | n.d. | NS | NS | NS |
| Homoeolog FC | NS | n.d. | n.d. | 1.4 | -0.9 | NS |

## Discussion

Allopolyploids face a complex array of challenges stemming both from whole genome duplication and from hybridization of divergent genomes. These challenges include maintaining stoichiometric balance among interacting molecules (Birchler and Veitia 2010, 2012, 2014, 2021), which may be even more problematic for interactions between the biparentally-inherited, organelle-targeted genes and those occurring in the maternally-coevolved organelles (Wolf and Hager 2006; Sharbrough *et al*. 2017). These potential cytonuclear incompatibilities may underlie observations of rapid and repeated return to single copy for organelle-targeted genes in polyploid species (De Smet *et al*. 2013; Li *et al*. 2016) and the expectation that any paternal cytonuclear homoeologs that exhibit deleterious interactions should evolve rapidly when not immediately lost (Rand *et al*. 2004; Sloan *et al*. 2014; Bock *et al*. 2014). Evidence from homoploid hybrids (Turelli and Moyle 2007; Greiner *et al*. 2011; Bock *et al*. 2014) suggests that stabilizing cytonuclear interactions is key to establishing a successful lineage, and surveys of Rubisco in diverse plant lineages (Gong *et al*. 2012, 2014) report differential homoeolog retention, biased expression, and asymmetric gene conversion favoring maternal homoeologs, although exceptions exist (Wang *et al*. 2017a; Zhai *et al*. 2019).

Emerging research into cytonuclear accommodation in allopolyploid species both supports and contradicts *a priori* cytonuclear expectations of maternal bias (Gong *et al*. 2012, 2014; Sehrish *et al*. 2015; Wang *et al*. 2017a, 2017b; Ferreira de Carvalho *et al*. 2019; Zhai *et al*. 2019; Shan *et al*. 2020; Sharbrough *et al*. 2021), meaning that only some allopolyploids exhibited maternal bias in some cytonuclear genes whereas others did not. Against this backdrop of observations and expectations, we surveyed global gene expression for five allopolyploid species and one synthetic representing four different genera encompassing a wide range of divergence times to evaluate the extent to which gene expression patterns change in accordance with cytonuclear expectations. While we analyze these data here for the purpose of evaluating gene expression changes in allopolyploids, we also note that because these data include ncRNAs and organellar reads, they represent a valuable resource for the allopolyploid community.

### Total gene expression exhibits limited evidence of cytonuclear maternal expression level dominance

Cytonuclear imbalance in polyploids could potentially arise due to the changes in dosage balance between organellar and nuclear genomes that accompany polyploidy. In response, the nascent polyploid might be expected to experience selection to mitigate any dosage-related detrimental effects by altering total gene expression in either the organelle or nucleus. Changes in organelle copy number and/or genome copy per organelle have been associated with polyploidy (Bingham 1968; Beversdorf 1979; Dean and Leech 1982; Butterfass 1987; Murti *et al*. 2012; Oberprieler *et al*. 2019; Coate *et al*. 2020; He *et al*. 2021; Fernandes Gyorfy *et al*. 2021), and these have been associated with cytonuclear compensation at the expression level (Doyle and Coate 2019; Coate *et al*. 2020). On the other hand, it is common for polyploids to undergo rapid changes in nuclear expression (Chen 2007; Doyle *et al*. 2008; Freeling 2009; Gaeta and Pires 2010; Jackson and Chen 2010; Salmon *et al*. 2010; Grover *et al*. 2012; Madlung and Wendel 2013; Yoo *et al*. 2014; Song and Chen 2015; Bao *et al*. 2019; Gallagher *et al*. 2020), which could include changes that compensate for deleterious cytonuclear stoichiometric imbalances.

In the present study, we characterized how total expression of nuclear-encoded cytonuclear genes changes relative to the rest of the transcriptome and whether those changes are biased toward the maternal parent. We evaluated each polyploid for evidence of maternally-biased expression level dominance (ELD) in cytonuclear genes that is statistically different from any global, or background, bias exhibited by genes not involved in cytonuclear processes. Our expectation was that we would observe some degree of ELD for cytonuclear genes that might provide evidence of cytonuclear compensation to coordinate expression with the maternally co-evolved organelles. While three of the five polyploids (*i.e., Arachis hypogea*, *Gossypium hirsutum*, and *Gossypium barbadense*) exhibited a general bias toward maternal ELD, only *Arachis hypogea* and *Chenopodium quinoa* exhibited an excess of maternal ELD in cytonuclear genes (in general) relative to the remaining transcriptome (*i.e.,* NOT; Table 4), with only 1-2 categories exhibiting evidence of significant ELD (DNI in both species and MI in *Chenopodium quinoa*). Interestingly, however, *Gossypium barbadense*, while not exhibiting a general parental bias in cytonuclear ELD, did exhibit maternally-biased ELD in the PI cytonuclear category alone. While these results suggest that global ELD in cytonuclear genes is not a *general* consequence of cytonuclear accommodation, it is noteworthy that in many cases, and for all species, the number of genes exhibiting maternally-biased ELD in cytonuclear categories does exceed the expected number (although not significantly so). This may be a function of the limited numbers of genes in each category, trends of partial yet non-ubiquitous maternally-biased ELD in cytonuclear categories, and/or both. While we also evaluated patterns of transgressive expression in cytonuclear categories relative to non-organelle-targeted genes, we did not find evidence of biased transgressive expression that would indicate a global up- or down-regulation of cytonuclear genes to compensate for the number of organelles/organelle genomes; however, we again note that most categories were limited in membership, leading to low statistical power.

### Variability in cytonuclear homoeolog expression patterns

Cytonuclear imbalance in allopolyploid species can also arise from incompatibilities between the organellar genomes and the more divergent paternal cytonuclear genes, and we expect these to become more common as the divergence time between progenitor genomes increases.

Reconciliation of potentially maladaptive mutations is possible through a variety of mechanisms, as previously noted (Gong *et al*. 2012, 2014; Sharbrough *et al*. 2017). For example, at the genomic level, gene loss and maternally-biased gene conversion could either remove or “correct” maladaptive mutations acquired by the paternal genome since its divergence from a common ancestor, minimizing their deleterious potential (Sharbrough *et al*. 2021).

With respect to expression, compensation for maladaptive paternal mutations could present as a combination of up-regulated maternal homoeologs and/or down-regulated paternal homoeologs. This, however, does not appear to be a global reaction to allopolyploidy in the species surveyed. When we compared homoeolog expression for each of the six allopolyploid species with their respective diploid progenitor genomes, we observed no clear and consistent pattern of homoeolog up-/down-regulation within polyploids and/or for any of the cytonuclear categories. At most, any given polyploid displayed two cytonuclear categories consistent with our biological expectations of excess maternal up-regulation and/or paternal down-regulation (Figure 1), and concomitantly have as many or more categories that directly contradict our cytonuclear predictions (*i.e.,* enrichment of maternal down-regulation and/or paternal up-regulation).

Individual cytonuclear categories were no more consistent, with the MI category being most frequently consistent (*i.e.,* agreed with expectations in two species, *Arabidopsis suecica* and *Arachis hypogea*), while also being contradictory in the same number of species (*Arachis* IpaDur1 and *Chenopodium quinoa*).

Maternal homoeolog expression bias (*i.e.,* genes where maternal expression outweighs paternal, irrespective of diploid expression) was similarly intermittent in cytonuclear categories. When compared to any global HEB exhibited by each species, few cytonuclear categories exhibited an excess of maternal HEB (*i.e., Arabidopsis suecica* PI/PNI and *Gossypium hirsutum* DI only; Table 5). Interestingly, a single category in *Chenopodium quinoa* (MI) and several in *Gossypium barbadense* (DNI/PI/PNI) exhibited an excess of *paternal* HEB, which is contrary to cytonuclear expectations. We do note, however, that these relative expression biases are often parentally inherited, as noted by the previous analysis.

Importantly, our analytical methodology was designed to disentangle parental or progenitor legacy effects (*i.e.*, differences at the diploid level vertically inherited in the polyploids at formation) from evolved cytonuclear responses subsequent to polyploid formation. When we combined our assessment of homoeolog expression differences with legacy parental effects (Table 7), we find that not only do targeting (*i.e.*, cytonuclear category) and legacy diploid expression influence the difference in homoeolog expression, but there is also an interactive effect between targeting and legacy expression differences. Interestingly, however, many of the fixed effects are not congruent with our expectations under the cytonuclear hypotheses, *i.e.*, that the difference in *rlog* counts between maternal and paternal homoeologs should be greater in the cytonuclear categories (or positive relative to the intercept established by NOT genes). Contrasts between each cytonuclear category and NOT genes also exhibited sporadic significance and were frequently incongruent with expectations (*i.e.,* that the cytonuclear categories would exhibit greater HEB when accounting for diploid legacy) for most species. Only *Arabidopsis suecica* showed significant, congruent cytonuclear effects for most categories, suggesting that DNI, MI/MNI, and PNI were generally composed of genes whose maternal HEB was greater than expected by NOT and diploid legacy.

In light of previous research that both supports and contrasts the results presented here, we speculate that cytonuclear accommodation is variable among lineages, among cytonuclear categories, and among genes within categories themselves. It also may be that for most genes (and especially those in the organellar genomes, which experience low mutation rates), the rates of molecular evolution are too low to permit signals of cytonuclear selection to become evident on the divergence scales studied here. It is possible, for example, that cytonuclear selection is ongoing and even pervasive, but that for the most part it is subtle, involving expression level changes or genomic signatures that simply do not rise to the level of statistical significance given the timescales encompassed by the allopolyploids studied here. Some polyploids, such as *Arabidopsis suecica*, provide a modest level of support for our *a priori* expectations for cytonuclear accommodation vis-á-vis gene expression, whereas others, such as *Gossypium barbadense*, contradict expectations more frequently than not. The variability in our observations may suggest that species with fewer cytonuclear-congruent expression changes either have fewer detrimental cytonuclear incompatibilities and/or have other methods for resolving deleterious conflict between the co-evolved maternal subgenome, the potentially detrimental paternal homoeologs, and the cytoplasmically inherited organelles.

## Data Availability Statement

All sequence data used in the analysis are available from NCBI under PRJNA726938, and all scripts used to analyze the data are available from Github under https://github.com/Wendellab/CytonuclearExpression commit XXXXXXXX.

## Acknowledgements

We thank Andreas Madlung and Roswitha Shmickl for providing *Arabidopsis* seeds, and David Brenner for providing *Chenopodium* plants. We thank Kenneth McCabe for supportive plant care in the Pohl Conservatory, and ResearchIT at Iowa State University for computational support. This work utilized the Summit supercomputer, which is supported by the National Science Foundation (awards ACI-1532235 and ACI-1532236), the University of Colorado Boulder, and Colorado State University. The Summit supercomputer is a joint effort of the University of Colorado Boulder and Colorado State University. This work was supported by the National Science Foundation (NSF) awards IOS-1829176 (DBS, JFW, JS, and CEG), IOS-2145811 (JS), and U.S. Department of Agriculture ARS 58-6066-0-066 (DGP, CEG, and JFW).

---

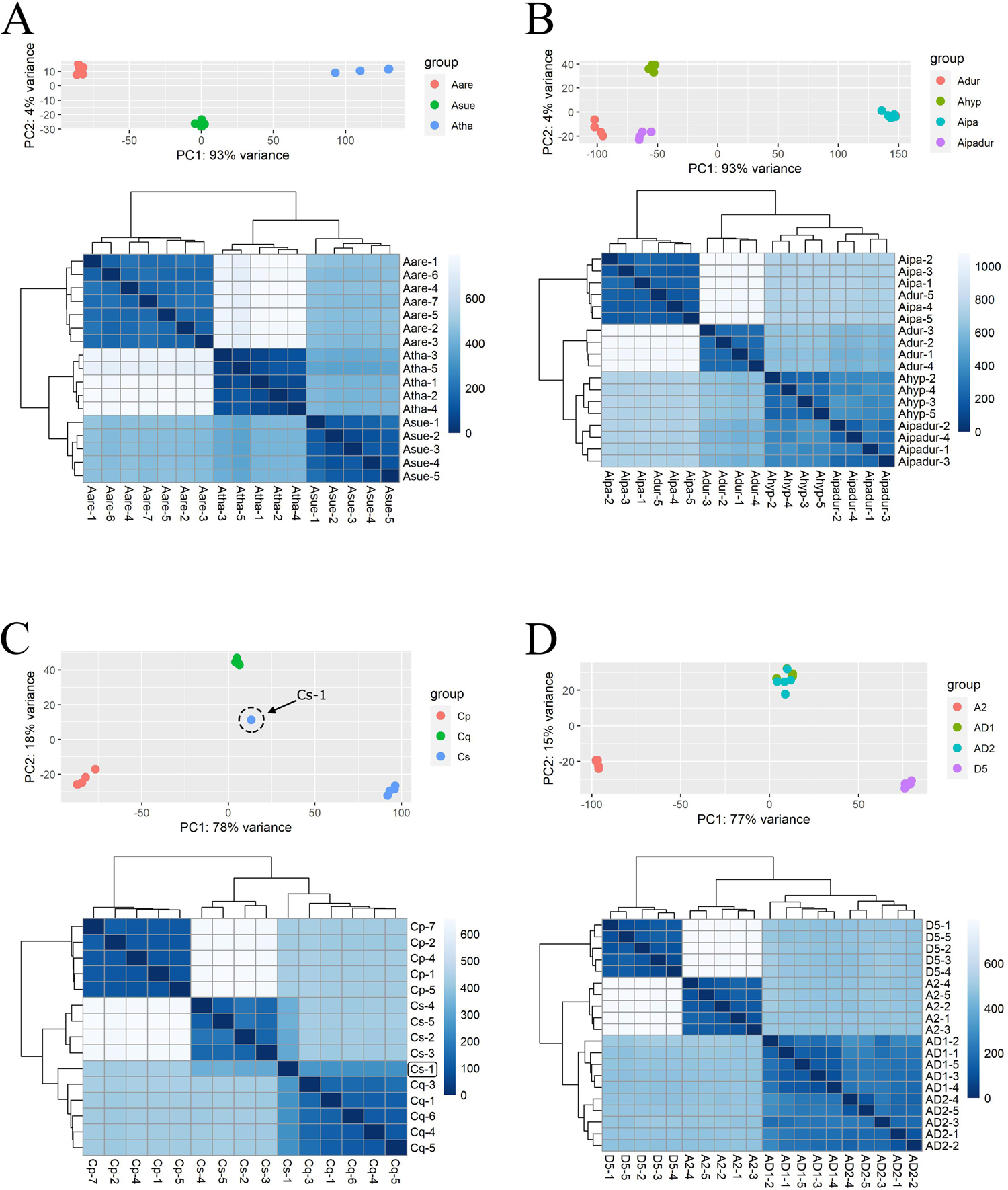

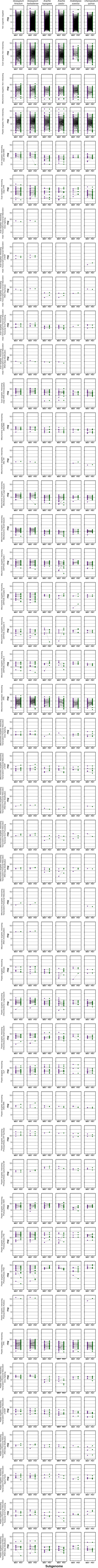

**Supplementary Table 1.** Species and accession used, with ploidy levels.

| Genus | Species | Accession | Ploidy |
| --- | --- | --- | --- |
| ***Chenopodium*** |  |  |  |
|  | *C. quinoa* | QQ74 | tetraploid |
|  | *C. pallidicaule* | PI 478407 | diploid |
|  | *C. suecicum* | Not Available | diploid |
| ***Gossypium*** |  |  |  |
|  | *G. hirsutum* | TM1 | tetraploid |
|  | *G. barbadense* | GB 3-79 | tetraploid |
|  | *G. arboreum* | A2 101 | diploid |
|  | *G. raimondii* | JFW | diploid |
| ***Arabidopsis*** |  |  |  |
|  | *A. suecica* | CS22505 | tetraploid |
|  | *A. thaliana* | Landsberg CS69111 | diploid |
|  | *A. arenosa* | 900118 | diploid |
| ***Arachis*** |  |  |  |
|  | *A. hypogea* | Tifrunner | tetraploid |
|  | *A. ipaensis* x *A. duranensis* | Bertioli et al. 2019 | tetraploid |
|  | *A. ipaensis* | GK30076 | diploid |
|  | *A. duranensis* | V14167 | diploid |

**Supplementary Table 2.** Genomic and transcriptomic references used in reference transcriptome curation.

|  |  |  |  |
| --- | --- | --- | --- |
| Genus | chloroplast | mitochondria | nuclear |
| *Arabidopsis* | NC_000932 | NC_037304 | Novikova et al. 2017 |
| *Arachis* | NC_037358 | NCBI SRA SRR14414925 and PeanutBase | Bertioli et al. 2019 |
| *Chenopodium* | MK159176 | MK182703 | Jarvis et al. 2017 |
| *Gossypium* | NC_007944 | JX065074 | Chen et al. 2020 |

**Supplementary Table 3.** Average sequencing and mapping results for RNA-seq libraries, by species.

|  | replicates | # fragments<br>sequenced (in<br>millions) | % fragments mapped | % fragments mapped<br>to chloroplast | % fragments mapped<br>to mitochondria |
| --- | --- | --- | --- | --- | --- |
| *Arabidopsis thaliana* | 5 | 65.6 (50.4 - 81.4) | 85% (83 - 87%) | 73% (69 - 75%) | 1% (1 - 2%) |
| *Arabidopsis suecica* | 5 | 55.1 (37.3 - 66.5) | 76% (67 - 81%) | 67% (63 - 70%) | 2% (2 - 3%) |
| *Arabidopsis arenosa* | 7 | 61.6 (53.5 - 83.0) | 66% (57 - 75%) | 66% (56 - 71%) | 3% (2 - 3%) |
| *Arachis duranensis* | 5 | 38.7 (33.1 - 43.3) | 82% (80 - 85%) | 67% (56 - 76%) | 1% (1 - 2%) |
| *Arachis ipaensis* | 5 | 40.5 (34.3 - 45.0) | 81% (80 - 82%) | 57% (53 - 59%) | 1% (1 - 2%) |
| *Arachis hypogea* | 4 | 31.3 (26.1 - 43.5) | 85% (83 - 86%) | 66% (58 - 74%) | 1% |
| *Arachis IpaDur1* | 4 | 33.9 (30.1 - 41.2) | 79% (75 - 83%) | 62% (56 - 71%) | 2% (1 - 2%) |
| *Chenopodium pallidicaule* | 5 | 44.5 (41.0 - 50.4) | 61% (55 - 68%) | 73% (72 - 75%) | 2% (2 - 3%) |
| *Chenopodium suecicum* | 4 | 41.2 (37.9 - 45.3) | 57% (40 - 69%) | 74% (73 - 76%) | 1% (1 - 2%) |
| *Chenopodium quinoa* | 5 | 47.0 (33.8 - 59.9) | 68% (56 - 80%) | 74% (72 - 76%) | 1% (1 - 2%) |
| *Gossypium arboreum* | 5 | 51.0 (44.6 - 57.7) | 74% (65 - 79%) | 56% (46 - 62%) | 2% |
| *Gossypium raimondii* | 5 | 68.7 (52.0 - 89.9) | 59% (50 - 67%) | 43% (38 - 50%) | 3% (2 - 4%) |
| *Gossypium hirsutum* | 5 | 59.8 (53.7 - 71.7) | 78% (67 - 85%) | 52% (47 - 58%) | 2% (2 - 3%) |
| *Gossypium barbadense* | 5 | 55.5 (49.3 - 68.5) | 79% (76 - 81%) | 53% (43 - 63%) | 2% (1 - 2%) |

**Supplementary Table 4.**
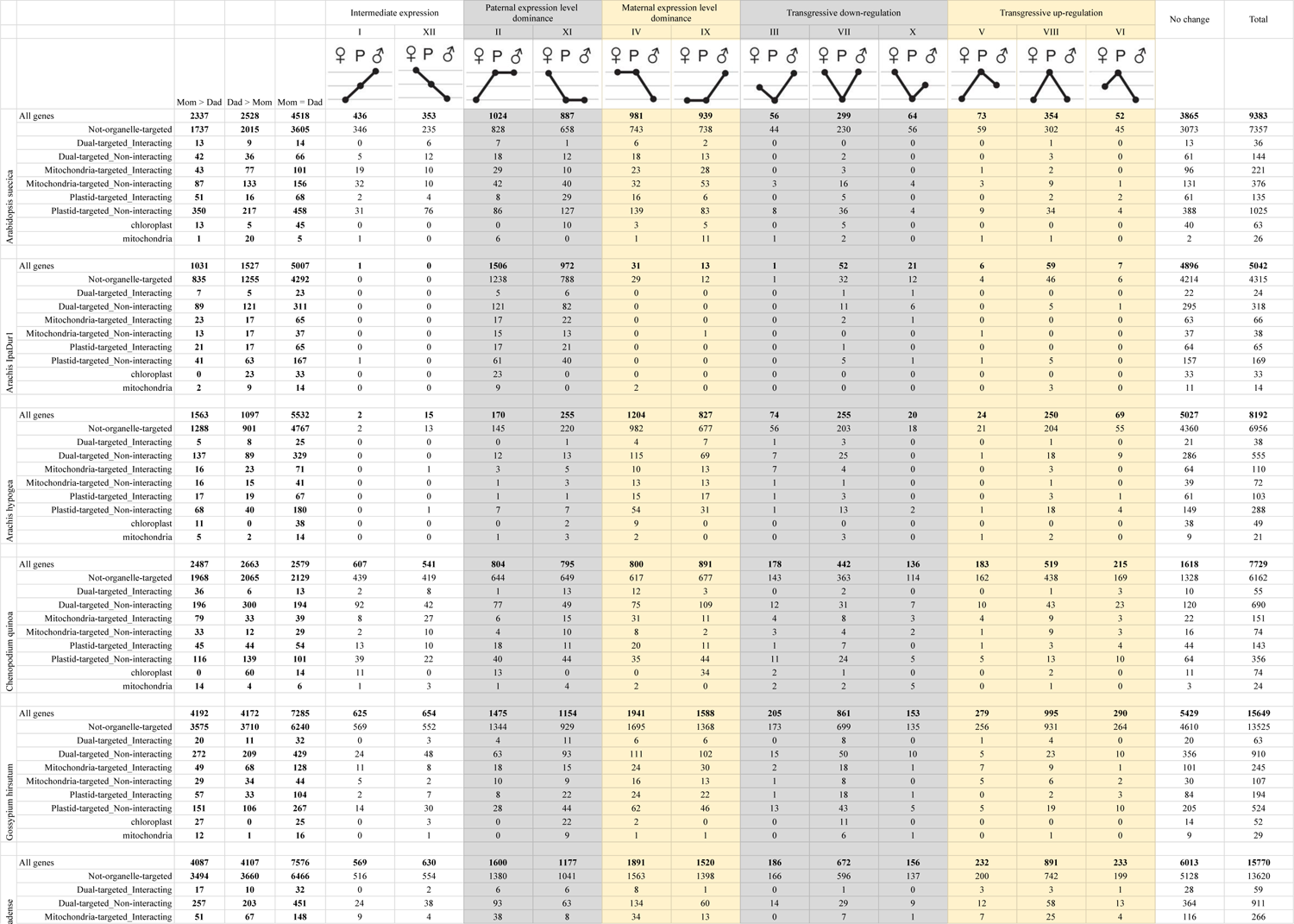
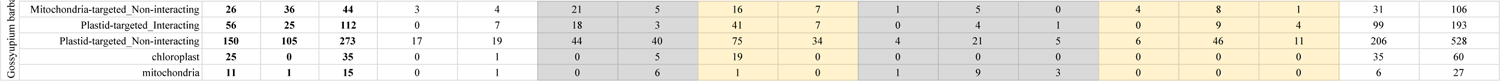
Expression level dominance in six allotetraploids.

**Supplementary Table 5.**
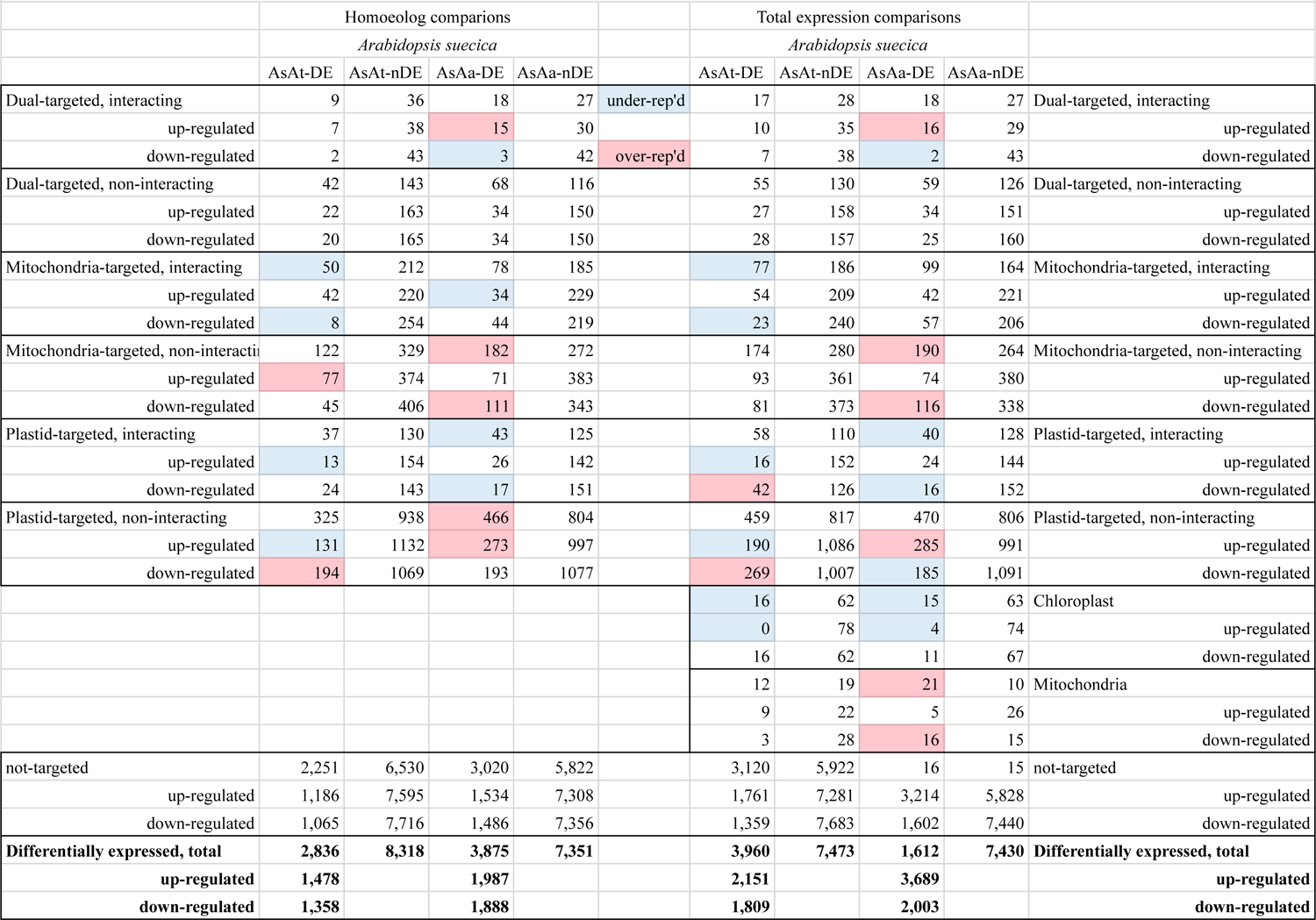
Genes exhibiting differential expression (DE) relative to those not exhibiting differential expression (nDE) relative to the diploid parents. The homoeolog-based comparison refers to DE between maternal parent-maternal homoeolog or the paternal parent-paternal homoeolog comparison. Total expression evalutes DE between the indicated diploid parent and the total gene expression in the polyploid (represented by the sum of both homoeologs). For *Arabidopsis*, AsAt denotes the maternal parent and AsAa denotes the paternal.

**Supplementary Table 6.**
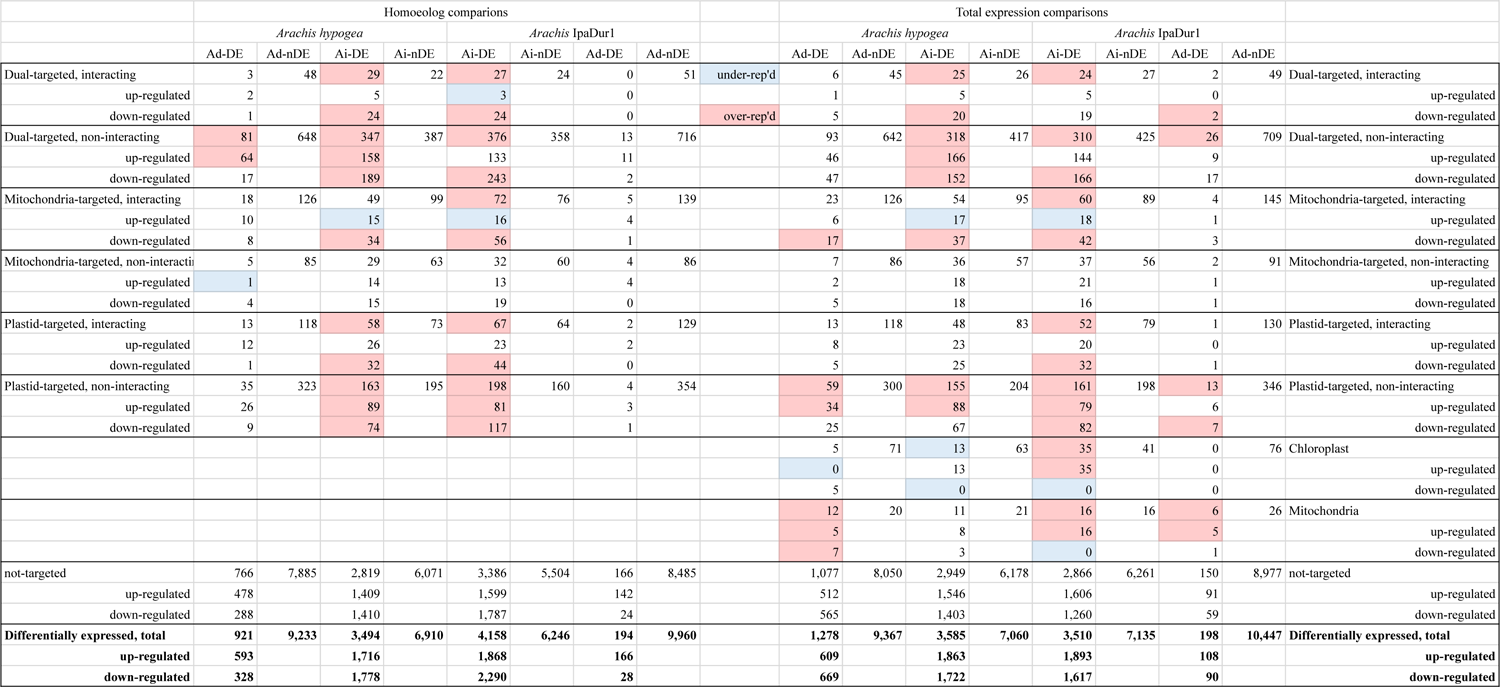
Genes exhibiting differential expression (DE) relative to those not exhibiting differential expression (nDE) relative to the diploid parents. The homoeolog-based comparison refers to DE between maternal parent-maternal homoeolog or the paternal parent-paternal homoeolog comparison. Total expression evalutes DE between the indicated diploid parent and the total gene expression in the polyploid (represented by the sum of both homoeologs). For *Arachis*, Ad denotes the *Arachis duranensis* (maternal to *A. hypogea* and paternal to IpaDur1) and Ai denotes *Arachis ipaensis* (maternal to IpaDur1 and paternal to *A. hypogea*).

**Supplementary Table 7.**
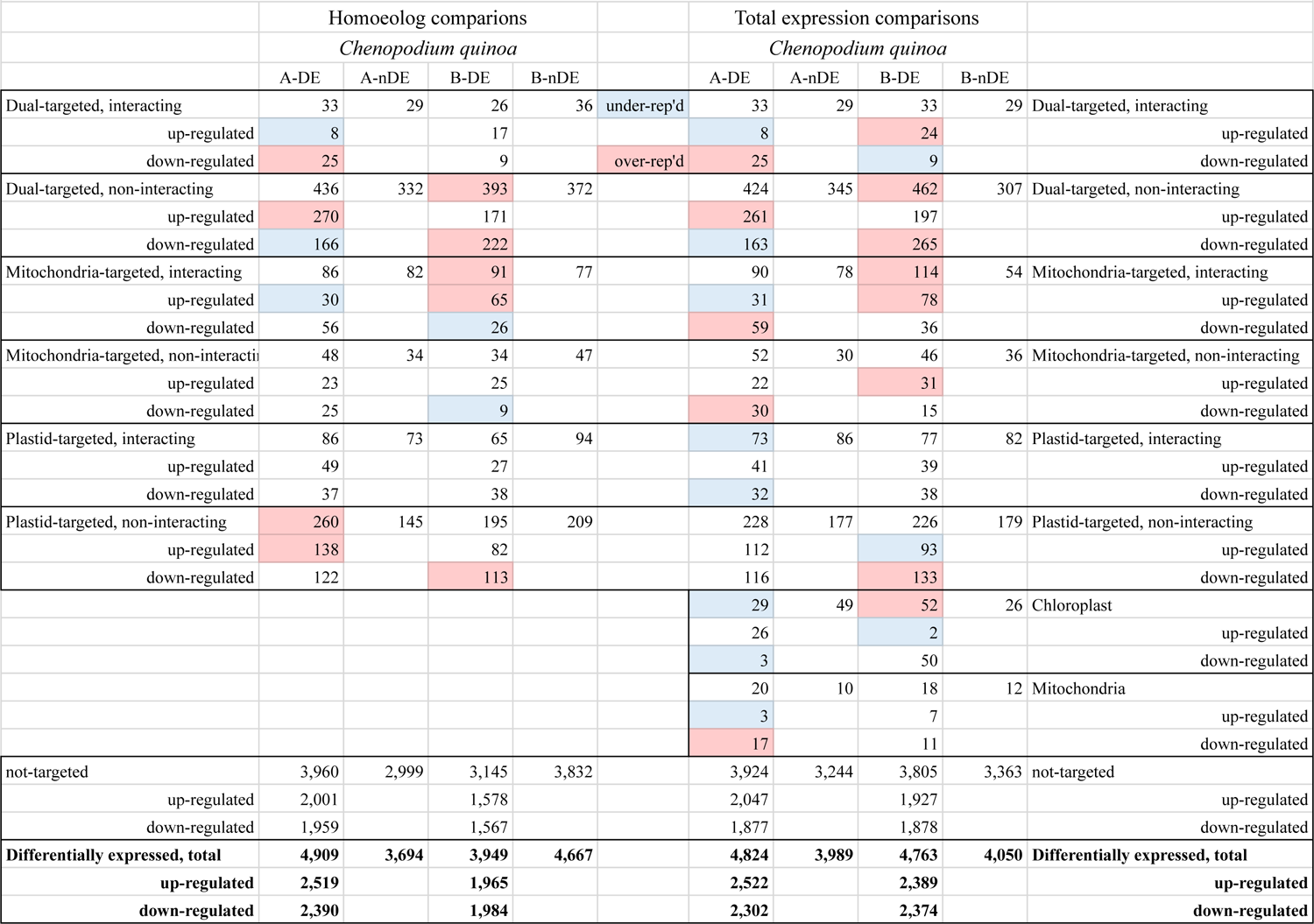
Genes exhibiting differential expression (DE) relative to those not exhibiting differential expression (nDE) relative to the diploid parents. The homoeolog-based comparison refers to DE between maternal parent-maternal homoeolog or the paternal parent-paternal homoeolog comparison. Total expression evalutes DE between the indicated diploid parent and the total gene expression in the polyploid (represented by the sum of both homoeologs). For *Chenopodium*, A denotes the maternal parent and B denotes the paternal.

|  | Homoeolog comparisons |  |  |  |  | Total expression comparisons |  |  |  |  |
| --- | --- | --- | --- | --- | --- | --- | --- | --- | --- | --- |
|  | *Chenopodium quinoa* |  |  |  |  | *Chenopodium quinoa* |  |  |  |  |
|  | A-DE | A-nDE | B-DE | B-nDE |  | A-DE | A-nDE | B-DE | B-nDE |  |
| Dual-targeted, interacting | 33 | 29 | 26 | 36 | under-rep'd | 33 | 29 | 33 | 29 | Dual-targeted, interacting |
| up-regulated | 8 |  | 17 |  |  | 8 |  | 24 |  | up-regulated |
| down-regulated | 25 |  | 9 |  | over-rep'd | 25 |  | 9 |  | down-regulated |
| Dual-targeted, non-interacting | 436 | 332 | 393 | 372 |  | 424 | 345 | 462 | 307 | Dual-targeted, non-interacting |
| up-regulated | 270 |  | 171 |  |  | 261 |  | 197 |  | up-regulated |
| down-regulated | 166 |  | 222 |  |  | 163 |  | 265 |  | down-regulated |
| Mitochondria-targeted, interacting | 86 | 82 | 91 | 77 |  | 90 | 78 | 114 | 54 | Mitochondria-targeted, interacting |
| up-regulated | 30 |  | 65 |  |  | 31 |  | 78 |  | up-regulated |
| down-regulated | 56 |  | 26 |  |  | 59 |  | 36 |  | down-regulated |
| Mitochondria-targeted, non-interacting | 48 | 34 | 34 | 47 |  | 52 | 30 | 46 | 36 | Mitochondria-targeted, non-interacting |
| up-regulated | 23 |  | 25 |  |  | 22 |  | 31 |  | up-regulated |
| down-regulated | 25 |  | 9 |  |  | 30 |  | 15 |  | down-regulated |
| Plastid-targeted, interacting | 86 | 73 | 65 | 94 |  | 73 | 86 | 77 | 82 | Plastid-targeted, interacting |
| up-regulated | 49 |  | 27 |  |  | 41 |  | 39 |  | up-regulated |
| down-regulated | 37 |  | 38 |  |  | 32 |  | 38 |  | down-regulated |
| Plastid-targeted, non-interacting | 260 | 145 | 195 | 209 |  | 228 | 177 | 226 | 179 | Plastid-targeted, non-interacting |
| up-regulated | 138 |  | 82 |  |  | 112 |  | 93 |  | up-regulated |
| down-regulated | 122 |  | 113 |  |  | 116 |  | 133 |  | down-regulated |
|  |  |  |  |  |  | 29 | 49 | 52 | 26 | Chloroplast |
|  |  |  |  |  |  | 26 |  | 2 |  | up-regulated |
|  |  |  |  |  |  | 3 |  | 50 |  | down-regulated |
|  |  |  |  |  |  | 20 | 10 | 18 | 12 | Mitochondria |
|  |  |  |  |  |  | 3 |  | 7 |  | up-regulated |
|  |  |  |  |  |  | 17 |  | 11 |  | down-regulated |
| not-targeted | 3,960 | 2,999 | 3,145 | 3,832 |  | 3,924 | 3,244 | 3,805 | 3,363 | not-targeted |
| up-regulated | 2,001 |  | 1,578 |  |  | 2,047 |  | 1,927 |  | up-regulated |
| down-regulated | 1,959 |  | 1,567 |  |  | 1,877 |  | 1,878 |  | down-regulated |
| **Differentially expressed, total** | **4,909** | **3,694** | **3,949** | **4,667** |  | **4,824** | **3,989** | **4,763** | **4,050** | **Differentially expressed, total** |
| **up-regulated** | **2,519** |  | **1,965** |  |  | **2,522** |  | **2,389** |  | **up-regulated** |
| **down-regulated** | **2,390** |  | **1,984** |  |  | **2,302** |  | **2,374** |  | **down-regulated** |

**Supplementary Table 8.** Genes exhibiting differential expression (DE) relative to those not exhibiting differential expression (nDE) relative to the diploid parents. The homoeolog-based comparison refers to DE between maternal parent-matern

|  | Homoeolog comparisons |  |  |  |  |  |  |  |  | Total expression comparisons |  |  |  |  |  |  |  |  |
| --- | --- | --- | --- | --- | --- | --- | --- | --- | --- | --- | --- | --- | --- | --- | --- | --- | --- | --- |
|  | *Gossypium hirsutum* |  |  |  | *Gossypium barbadense* |  |  |  |  | *Gossypium hirsutum* |  |  |  | *Gossypium barbadense* |  |  |  |  |
|  | A-DE | A-nDE | D-DE | D-nDE | A-DE | A-nDE | D-DE | D-nDE |  | A-DE | A-nDE | D-DE | D-nDE | A-DE | A-nDE | D-DE | D-nDE |  |
| Dual-targeted, interacting | 22 | 54 | 33 | 43 | 25 | 51 | 28 | 48 | under-rep'd | 34 | 42 | 33 | 43 | 25 | 51 | 24 | 52 | Dual-targeted, interacting |
| up-regulated | 9 |  | 11 |  | 19 |  | 20 |  |  | 9 |  | 14 |  | 16 |  | 21 |  | up-regulated |
| down-regulated | 13 |  | 22 |  | 6 |  | 8 |  | over-rep'd | 25 |  | 19 |  | 9 |  | 3 |  | down-regulated |
| Dual-targeted, non-interacting | 327 | 758 | 486 | 602 | 336 | 749 | 450 | 638 |  | 390 | 701 | 461 | 630 | 397 | 694 | 437 | 654 | Dual-targeted, non-interacting |
| up-regulated | 108 |  | 236 |  | 167 |  | 297 |  |  | 144 |  | 221 |  | 234 |  | 291 |  | up-regulated |
| down-regulated | 219 |  | 250 |  | 169 |  | 153 |  |  | 246 |  | 240 |  | 163 |  | 146 |  | down-regulated |
| Mitochondria-targeted, interacting | 92 | 232 | 132 | 190 | 103 | 221 | 121 | 201 |  | 116 | 208 | 139 | 185 | 117 | 207 | 124 | 200 | Mitochondria-targeted, interacting |
| up-regulated | 43 |  | 57 |  | 85 |  | 92 |  |  | 51 |  | 56 |  | 96 |  | 91 |  | up-regulated |
| down-regulated | 49 |  | 75 |  | 18 |  | 29 |  |  | 65 |  | 83 |  | 21 |  | 33 |  | down-regulated |
| Mitochondria-targeted, non-interacting | 53 | 79 | 63 | 71 | 45 | 87 | 57 | 77 |  | 55 | 79 | 68 | 66 | 59 | 75 | 59 | 75 | Mitochondria-targeted, non-interacting |
| up-regulated | 30 |  | 33 |  | 27 |  | 35 |  |  | 31 |  | 34 |  | 41 |  | 36 |  | up-regulated |
| down-regulated | 23 |  | 30 |  | 18 |  | 22 |  |  | 24 |  | 34 |  | 18 |  | 23 |  | down-regulated |
| Plastid-targeted, interacting | 68 | 178 | 92 | 153 | 49 | 197 | 92 | 153 |  | 80 | 166 | 100 | 146 | 59 | 187 | 88 | 158 | Plastid-targeted, interacting |
| up-regulated | 17 |  | 47 |  | 25 |  | 77 |  |  | 20 |  | 44 |  | 40 |  | 74 |  | up-regulated |
| down-regulated | 51 |  | 45 |  | 24 |  | 15 |  |  | 60 |  | 56 |  | 19 |  | 14 |  | down-regulated |
| Plastid-targeted, non-interacting | 204 | 422 | 265 | 359 | 202 | 424 | 281 | 343 |  | 236 | 391 | 285 | 342 | 243 | 384 | 265 | 362 | Plastid-targeted, non-interacting |
| up-regulated | 63 |  | 132 |  | 103 |  | 192 |  |  | 83 |  | 144 |  | 143 |  | 178 |  | up-regulated |
| down-regulated | 141 |  | 133 |  | 99 |  | 89 |  |  | 153 |  | 141 |  | 100 |  | 87 |  | down-regulated |
|  |  |  |  |  |  |  |  |  |  | 50 | 27 | 16 | 61 | 6 | 71 | 24 | 53 | Chloroplast |
|  |  |  |  |  |  |  |  |  |  | 0 |  | 5 |  | 0 |  | 24 |  | up-regulated |
|  |  |  |  |  |  |  |  |  |  | 50 |  | 11 |  | 6 |  | 0 |  | down-regulated |
|  |  |  |  |  |  |  |  |  |  | 20 | 15 | 15 | 20 | 22 | 13 | 20 | 15 | Mitochondria |
|  |  |  |  |  |  |  |  |  |  | 1 |  | 4 |  | 0 |  | 5 |  | up-regulated |
|  |  |  |  |  |  |  |  |  |  | 19 |  | 11 |  | 22 |  | 15 |  | down-regulated |
| not-targeted | 5,203 | 10,505 | 7,231 | 8,635 | 4,765 | 10,943 | 6,484 | 9,382 |  | 6,561 | 9,733 | 7,617 | 8,677 | 6,177 | 10,117 | 6,883 | 9,411 | not-targeted |
| up-regulated | 2,896 |  | 3,874 |  | 2,482 |  | 3,339 |  |  | 3,747 |  | 4,178 |  | 3,374 |  | 3,660 |  | up-regulated |
| down-regulated | 2,307 |  | 3,357 |  | 2,283 |  | 3,145 |  |  | 2,814 |  | 3,439 |  | 2,803 |  | 3,223 |  | down-regulated |
| Differentially expressed, total | 5,969 | 12,228 | 8,302 | 10,053 | 5,525 | 12,672 | 7,513 | 10,842 |  | 7,472 | 11,320 | 8,703 | 10,089 | 7,077 | 11,715 | 7,880 | 10,912 | Differentially expressed, total |
| up-regulated | 3,166 |  | 4,390 |  | 2,908 |  | 4,052 |  |  | 4,085 |  | 4,691 |  | 3,944 |  | 4,351 |  | up-regulated |
| down-regulated | 2,803 |  | 3,912 |  | 2,617 |  | 3,461 |  |  | 3,387 |  | 4,012 |  | 3,133 |  | 3,529 |  | down-regulated |

**Supplementary Table 9.**
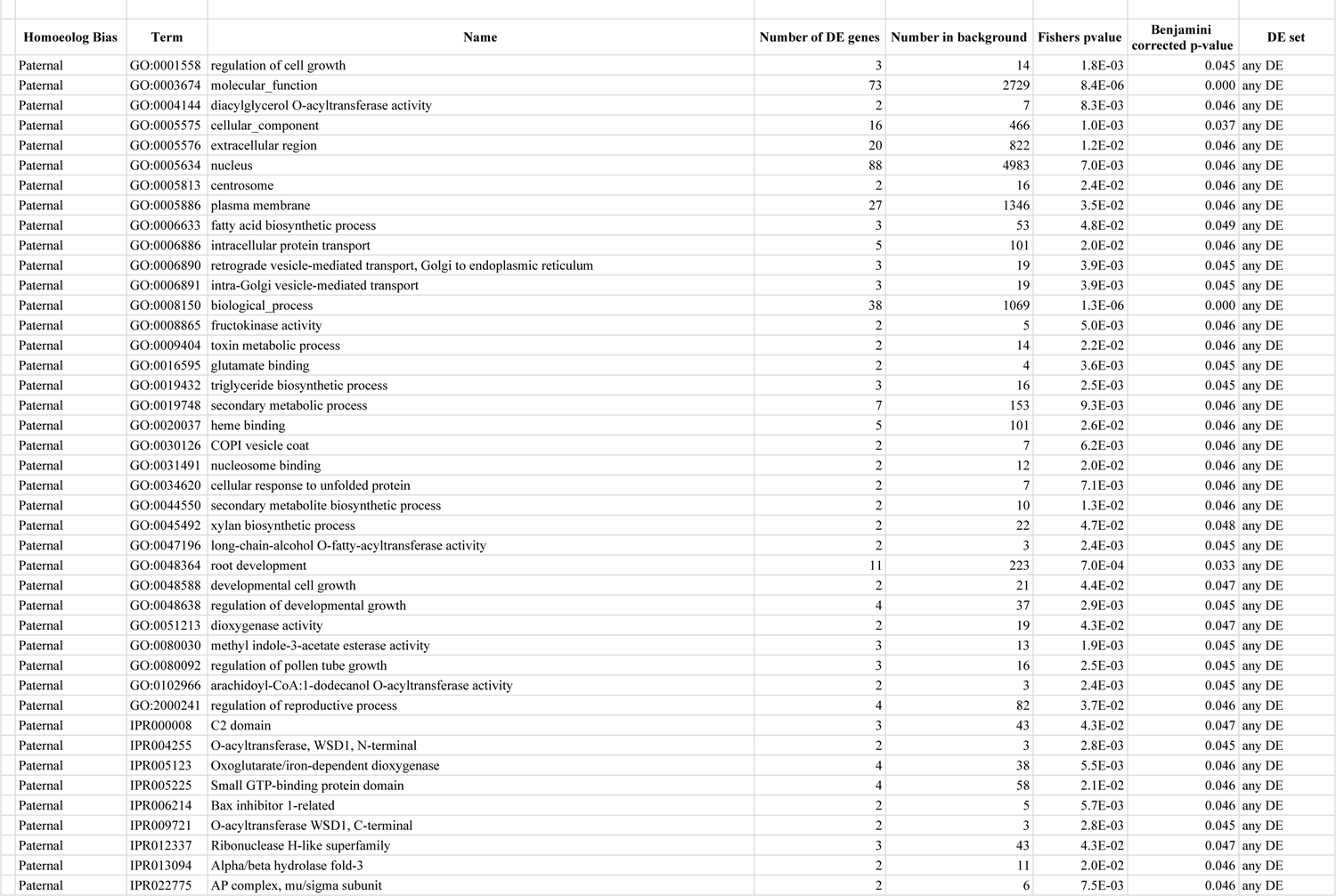
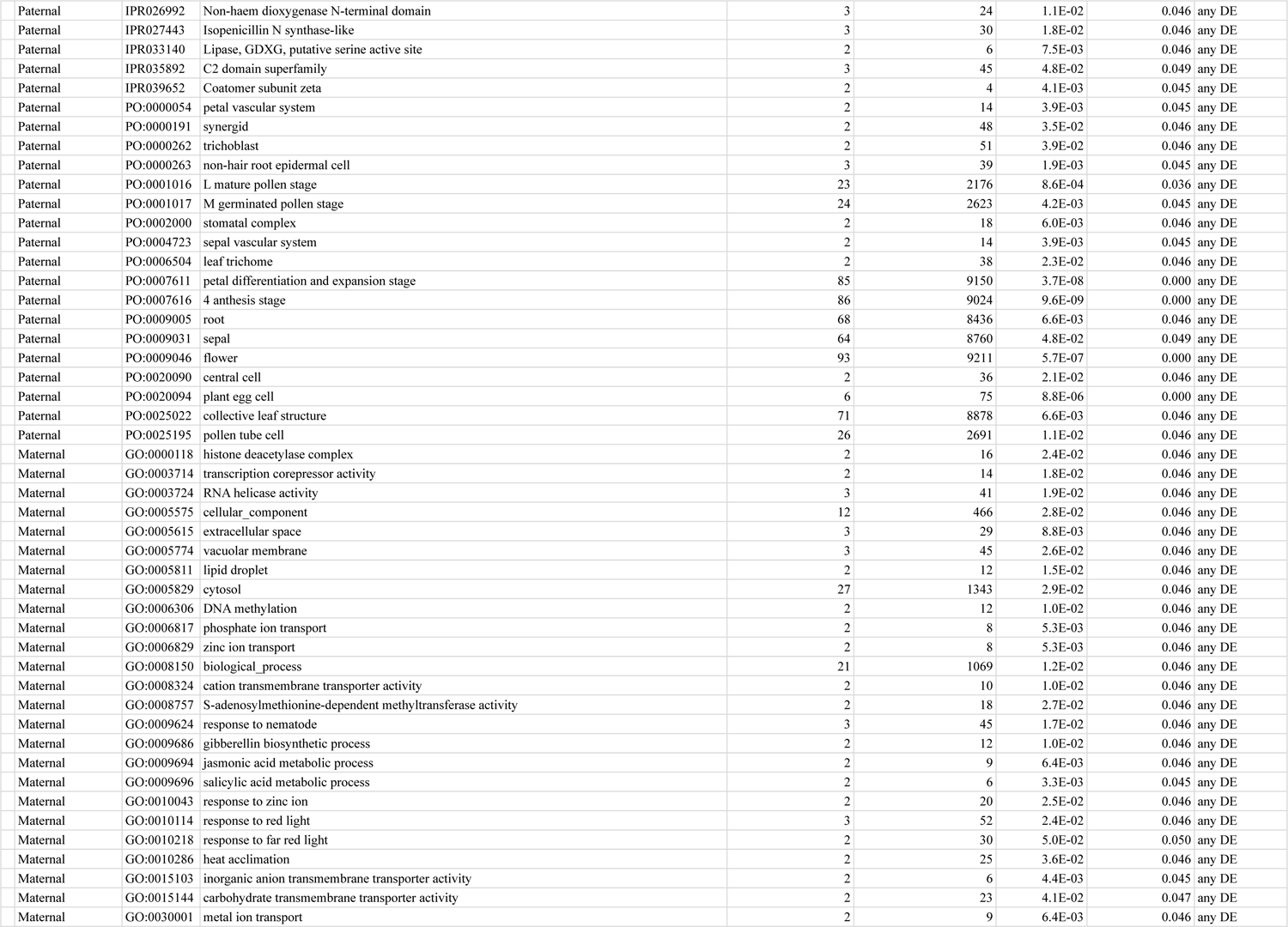
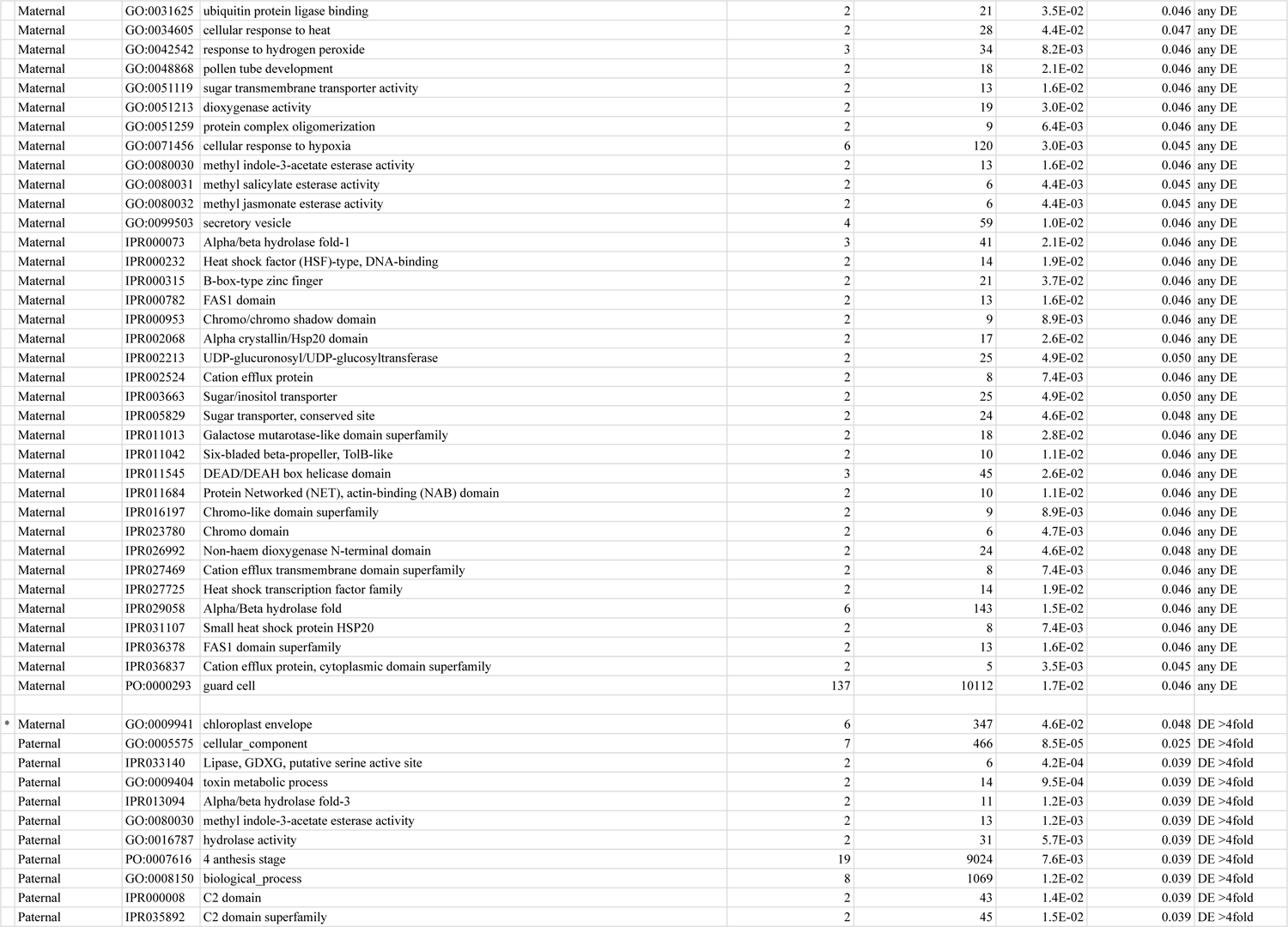
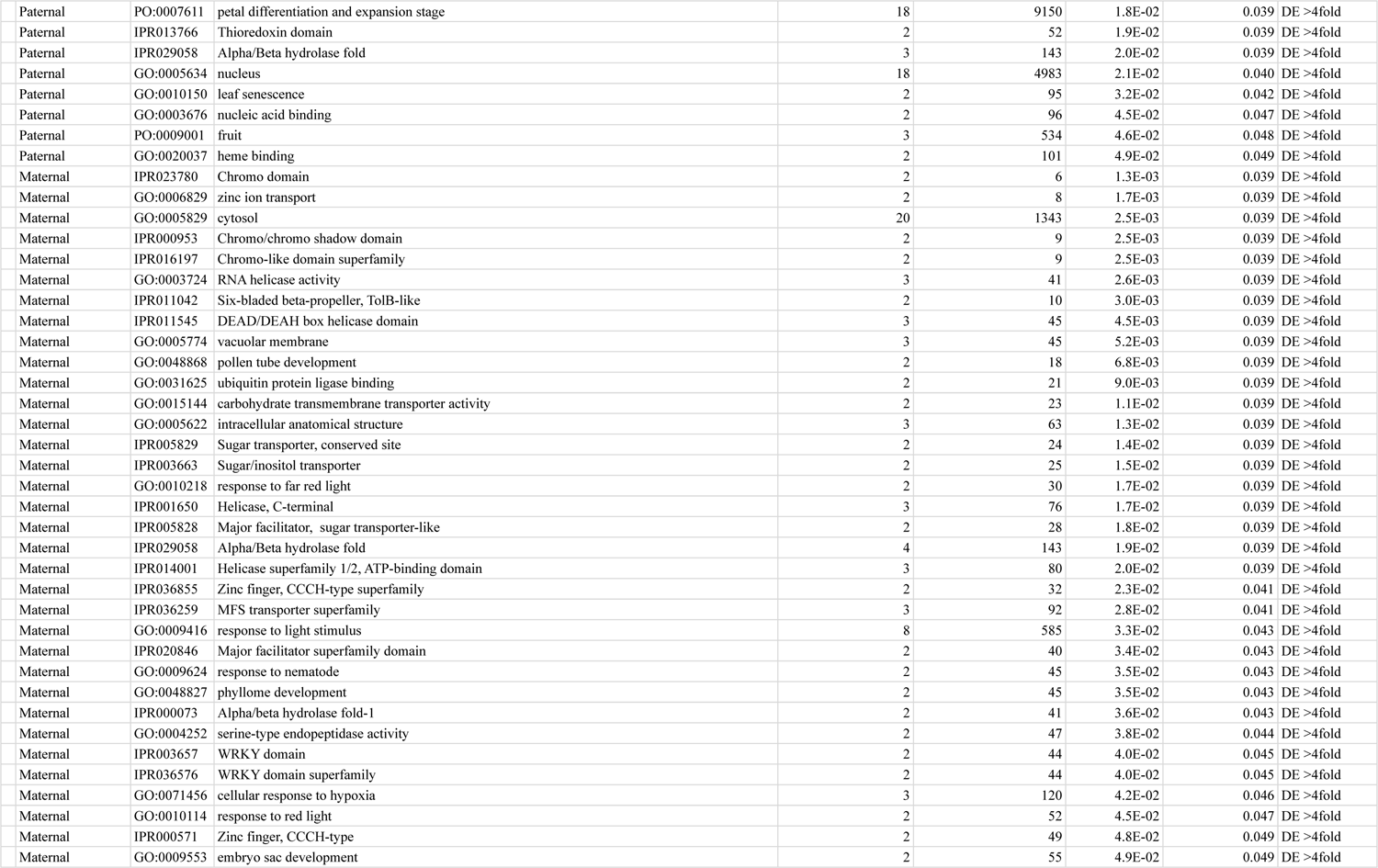
Enriched ontology terms (Benjamini corrected p-value < 0.05) for modules comprised of differentially expressed homoeologs with greater than fourfold difference in expression. The homoeolog bias columns indicates general bias for that functional module. The bottom half of the table contains enriched terms from the set of DE genes whose difference in fold-change between the subgenomes is greater than fourfold compared to the diploid genomes. Modules containing only one DE gene were excluded as unreliable.

**Supplementary Table 10.** Enriched ontology terms (Benjamini corrected p-value < 0.05) for modules comprised of differentially expressed homoeologs with greater than fourfold difference in expression. The bottom half of the table contains enriched terms from the set of DE genes whose difference in fold-change between the subgenomes is greater than fourfold compared to the diploid genomes. Modules containing only one DE gene were excluded as unreliable.

|  |  |  |  |  |  |  |  |  |
| --- | --- | --- | --- | --- | --- | --- | --- | --- |
| Species | Homoeolog Bias | Term | Name | Number of DE genes | Number in background | Fishers p-value | Benjamini corrected p-value | DE set |
| *G. hirsutum* | Paternal bias | GO:0005634 | nucleus | 20 | 410 | 0.001 | 0.046 | any DE |
| *G. hirsutum* | Paternal bias | IPR006689 | Small GTPase superfamily, ARF/SAR type | 4 | 25 | 0.003 | 0.046 | any DE |
| *G. hirsutum* | Paternal bias | GO:0006357 | regulation of transcription by RNA polymerase II | 3 | 15 | 0.005 | 0.046 | any DE |
| *G. hirsutum* | Paternal bias | GO:0032784 | regulation of DNA-templated transcription, elongation | 2 | 6 | 0.01 | 0.046 | any DE |
| *G. hirsutum* | Paternal bias | IPR020904 | Short-chain dehydrogenase/reductase, conserved site | 3 | 21 | 0.012 | 0.046 | any DE |
| *G. hirsutum* | Paternal bias | IPR002119 | Histone H2A | 2 | 7 | 0.014 | 0.046 | any DE |
| *G. hirsutum* | Paternal bias | IPR032454 | Histone H2A, C-terminal domain | 2 | 7 | 0.014 | 0.046 | any DE |
| *G. hirsutum* | Paternal bias | GO:0046982 | protein heterodimerization activity | 4 | 48 | 0.021 | 0.046 | any DE |
| *G. hirsutum* | Paternal bias | IPR009072 | Histone-fold | 4 | 48 | 0.021 | 0.046 | any DE |
| *G. hirsutum* | Paternal bias | IPR023214 | HAD superfamily | 6 | 100 | 0.022 | 0.046 | any DE |
| *G. hirsutum* | Paternal bias | GO:0006508 | proteolysis | 10 | 237 | 0.022 | 0.046 | any DE |
| *G. hirsutum* | Paternal bias | GO:0003684 | damaged DNA binding | 2 | 10 | 0.024 | 0.046 | any DE |
| *G. hirsutum* | Paternal bias | GO:0005992 | trehalose biosynthetic process | 2 | 11 | 0.025 | 0.046 | any DE |
| *G. hirsutum* | Paternal bias | GO:0006289 | nucleotide-excision repair | 2 | 11 | 0.025 | 0.046 | any DE |
| *G. hirsutum* | Paternal bias | IPR001965 | Zinc finger, PHD-type | 5 | 77 | 0.026 | 0.046 | any DE |
| *G. hirsutum* | Paternal bias | IPR003337 | Trehalose-phosphatase | 2 | 11 | 0.028 | 0.046 | any DE |
| *G. hirsutum* | Paternal bias | IPR006379 | HAD-superfamily hydrolase, subfamily IIB | 2 | 11 | 0.028 | 0.046 | any DE |
| *G. hirsutum* | Paternal bias | IPR014756 | Immunoglobulin E-set | 3 | 30 | 0.029 | 0.046 | any DE |
| *G. hirsutum* | Paternal bias | GO:0007049 | cell cycle | 2 | 13 | 0.033 | 0.046 | any DE |
| *G. hirsutum* | Paternal bias | GO:0003677 | DNA binding | 27 | 895 | 0.036 | 0.046 | any DE |
| *G. hirsutum* | Paternal bias | GO:0000786 | nucleosome | 3 | 32 | 0.046 | 0.046 | any DE |
| *G. hirsutum* | Paternal bias | GO:0004252 | serine-type endopeptidase activity | 4 | 64 | 0.049 | 0.049 | any DE |
| *G. hirsutum* | Maternal bias | IPR014001 | Helicase superfamily 1/2, ATP-binding domain | 6 | 64 | 0.001 | 0.046 | any DE |
| *G. hirsutum* | Maternal bias | IPR001650 | Helicase, C-terminal | 6 | 65 | 0.001 | 0.046 | any DE |
| *G. hirsutum* | Maternal bias | IPR019821 | Kinesin motor domain, conserved site | 4 | 28 | 0.001 | 0.046 | any DE |
| *G. hirsutum* | Maternal bias | IPR008847 | Suppressor of forked | 2 | 3 | 0.002 | 0.046 | any DE |
| *G. hirsutum* | Maternal bias | GO:0005871 | kinesin complex | 4 | 41 | 0.002 | 0.046 | any DE |
| *G. hirsutum* | Maternal bias | IPR000629 | ATP-dependent RNA helicase DEAD-box, conserved site | 3 | 15 | 0.003 | 0.046 | any DE |
| *G. hirsutum* | Maternal bias | GO:0051082 | unfolded protein binding | 5 | 56 | 0.004 | 0.046 | any DE |
| *G. hirsutum* | Maternal bias | GO:0007018 | microtubule-based movement | 4 | 44 | 0.004 | 0.046 | any DE |
| *G. hirsutum* | Maternal bias | IPR001752 | Kinesin motor domain | 4 | 41 | 0.005 | 0.046 | any DE |
| *G. hirsutum* | Maternal bias | IPR011545 | DEAD/DEAH box helicase domain | 4 | 41 | 0.005 | 0.046 | any DE |
| *G. hirsutum* | Maternal bias | IPR027640 | Kinesin-like protein | 4 | 41 | 0.005 | 0.046 | any DE |
| *G. hirsutum* | Maternal bias | IPR002194 | Chaperonin TCP-1, conserved site | 2 | 6 | 0.006 | 0.046 | any DE |
| *G. hirsutum* | Maternal bias | IPR017998 | Chaperone tailless complex polypeptide 1 (TCP-1) | 2 | 6 | 0.006 | 0.046 | any DE |
| *G. hirsutum* | Maternal bias | IPR008480 | Protein of unknown function DUF761, plant | 3 | 23 | 0.007 | 0.046 | any DE |
| *G. hirsutum* | Maternal bias | IPR027410 | TCP-1-like chaperonin intermediate domain superfamily | 2 | 7 | 0.008 | 0.046 | any DE |
| *G. hirsutum* | Maternal bias | GO:0009116 | nucleoside metabolic process | 2 | 8 | 0.008 | 0.046 | any DE |
| *G. hirsutum* | Maternal bias | IPR025836 | Zinc knuckle CX2CX4HX4C | 3 | 24 | 0.008 | 0.046 | any DE |
| *G. hirsutum* | Maternal bias | IPR008972 | Cupredoxin | 5 | 76 | 0.008 | 0.046 | any DE |
| *G. hirsutum* | Maternal bias | GO:0003777 | microtubule motor activity | 4 | 44 | 0.009 | 0.046 | any DE |
| *G. hirsutum* | Maternal bias | IPR002022 | Pectate lyase | 3 | 26 | 0.01 | 0.046 | any DE |
| *G. hirsutum* | Maternal bias | IPR014014 | RNA helicase, DEAD-box type, Q motif | 3 | 26 | 0.01 | 0.046 | any DE |
| *G. hirsutum* | Maternal bias | IPR018082 | AmbAllergen | 3 | 26 | 0.01 | 0.046 | any DE |
| *G. hirsutum* | Maternal bias | IPR025558 | Domain of unknown function DUF4283 | 3 | 28 | 0.012 | 0.046 | any DE |
| *G. hirsutum* | Maternal bias | IPR000198 | Rho GTPase-activating protein domain | 2 | 9 | 0.012 | 0.046 | any DE |
| *G. hirsutum* | Maternal bias | IPR003107 | HAT (Half-A-TPR) repeat | 2 | 9 | 0.012 | 0.046 | any DE |
| *G. hirsutum* | Maternal bias | IPR008936 | Rho GTPase activation protein | 2 | 9 | 0.012 | 0.046 | any DE |
| *G. hirsutum* | Maternal bias | IPR003653 | Ulp1 protease family, C-terminal catalytic domain | 2 | 10 | 0.014 | 0.046 | any DE |
| *G. hirsutum* | Maternal bias | GO:0008017 | microtubule binding | 4 | 53 | 0.016 | 0.046 | any DE |
| *G. hirsutum* | Maternal bias | IPR007524 | Pectate lyase, N-terminal | 2 | 11 | 0.017 | 0.046 | any DE |
| *G. hirsutum* | Maternal bias | IPR002173 | Carbohydrate/purine kinase, PfkB, conserved site | 2 | 12 | 0.019 | 0.046 | any DE |
| *G. hirsutum* | Maternal bias | IPR002453 | Beta tubulin | 2 | 12 | 0.019 | 0.046 | any DE |
| *G. hirsutum* | Maternal bias | GO:0030570 | pectate lyase activity | 2 | 11 | 0.02 | 0.046 | any DE |
| *G. hirsutum* | Maternal bias | IPR000095 | CRIB domain | 2 | 13 | 0.022 | 0.046 | any DE |
| *G. hirsutum* | Maternal bias | IPR013838 | Beta tubulin, autoregulation binding site | 2 | 13 | 0.022 | 0.046 | any DE |
| *G. hirsutum* | Maternal bias | GO:0006396 | RNA processing | 3 | 41 | 0.023 | 0.046 | any DE |
| *G. hirsutum* | Maternal bias | IPR027413 | GroEL-like equatorial domain superfamily | 2 | 15 | 0.028 | 0.046 | any DE |
| *G. hirsutum* | Maternal bias | IPR029044 | Nucleotide-diphospho-sugar transferases | 5 | 105 | 0.028 | 0.046 | any DE |
| *G. hirsutum* | Maternal bias | GO:0006397 | mRNA processing | 2 | 17 | 0.028 | 0.046 | any DE |
| *G. hirsutum* | Maternal bias | IPR011707 | Multicopper oxidase, type 3 | 3 | 41 | 0.03 | 0.046 | any DE |
| *G. hirsutum* | Maternal bias | IPR011611 | Carbohydrate kinase PfkB | 2 | 16 | 0.031 | 0.046 | any DE |
| *G. hirsutum* | Maternal bias | IPR001117 | Multicopper oxidase, type 1 | 3 | 42 | 0.032 | 0.046 | any DE |
| *G. hirsutum* | Maternal bias | IPR027409 | GroEL-like apical domain superfamily | 2 | 17 | 0.034 | 0.046 | any DE |
| *G. hirsutum* | Maternal bias | IPR029056 | Ribokinase-like | 2 | 17 | 0.034 | 0.046 | any DE |
| *G. hirsutum* | Maternal bias | IPR001296 | Glycosyl transferase, family 1 | 2 | 18 | 0.038 | 0.046 | any DE |
| *G. hirsutum* | Maternal bias | IPR001305 | Heat shock protein DnaJ, cysteine-rich domain | 2 | 18 | 0.038 | 0.046 | any DE |
| *G. hirsutum* | Maternal bias | IPR002423 | Chaperonin Cpn60/TCP-1 family | 2 | 18 | 0.038 | 0.046 | any DE |
| *G. hirsutum* | Maternal bias | IPR024709 | Putative O-fucosyltransferase, plant | 2 | 18 | 0.038 | 0.046 | any DE |
| *G. hirsutum* | Maternal bias | IPR027356 | NPH3 domain | 2 | 18 | 0.038 | 0.046 | any DE |
| *G. hirsutum* | Maternal bias | GO:0006950 | response to stress | 3 | 52 | 0.041 | 0.046 | any DE |
| *G. hirsutum* | Maternal bias | IPR017975 | Tubulin, conserved site | 2 | 19 | 0.041 | 0.046 | any DE |
| *G. hirsutum* | Maternal bias | GO:0006457 | protein folding | 4 | 92 | 0.044 | 0.046 | any DE |
| *G. hirsutum* | Maternal bias | IPR000330 | SNF2-related, N-terminal domain | 2 | 20 | 0.045 | 0.046 | any DE |
| *G. hirsutum* | Maternal bias | GO:0031072 | heat shock protein binding | 2 | 18 | 0.045 | 0.046 | any DE |
| *G. hirsutum* | Paternal bias | IPR006689 | Small GTPase superfamily, ARF/SAR type | 3 | 25 | 0.001 | 0.037 | DE >4fold |
| *G. hirsutum* | Paternal bias | IPR023214 | HAD superfamily | 4 | 100 | 0.006 | 0.037 | DE >4fold |
| *G. hirsutum* | Paternal bias | GO:0006357 | regulation of transcription by RNA polymerase II | 2 | 15 | 0.008 | 0.037 | DE >4fold |
| *G. hirsutum* | Paternal bias | IPR002659 | Glycosyl transferase, family 31 | 2 | 19 | 0.009 | 0.037 | DE >4fold |
| *G. hirsutum* | Paternal bias | GO:0008378 | galactosyltransferase activity | 2 | 19 | 0.012 | 0.037 | DE >4fold |
| *G. hirsutum* | Paternal bias | GO:0006260 | DNA replication | 2 | 27 | 0.022 | 0.037 | DE >4fold |
| *G. hirsutum* | Paternal bias | IPR032675 | Leucine-rich repeat domain superfamily | 8 | 489 | 0.024 | 0.037 | DE >4fold |
| *G. hirsutum* | Paternal bias | IPR026992 | Non-haem dioxygenase N-terminal domain | 3 | 86 | 0.024 | 0.037 | DE >4fold |
| *G. hirsutum* | Paternal bias | GO:0006486 | protein glycosylation | 2 | 30 | 0.026 | 0.038 | DE >4fold |
| *G. hirsutum* | Paternal bias | IPR027443 | Isopenicillin N synthase-like | 3 | 95 | 0.031 | 0.04 | DE >4fold |
| *G. hirsutum* | Paternal bias | IPR005123 | Oxoglutarate/iron-dependent dioxygenase | 3 | 105 | 0.04 | 0.044 | DE >4fold |
| *G. hirsutum* | Paternal bias | GO:0006351 | transcription, DNA-templated | 3 | 93 | 0.04 | 0.044 | DE >4fold |
| *G. hirsutum* | Maternal bias | IPR002194 | Chaperonin TCP-1, conserved site | 2 | 6 | 0.001 | 0.037 | DE >4fold |
| *G. hirsutum* | Maternal bias | IPR017998 | Chaperone tailless complex polypeptide 1 (TCP-1) | 2 | 6 | 0.001 | 0.037 | DE >4fold |
| *G. hirsutum* | Maternal bias | IPR027410 | TCP-1-like chaperonin intermediate domain superfamily | 2 | 7 | 0.001 | 0.037 | DE >4fold |
| *G. hirsutum* | Maternal bias | GO:0030570 | pectate lyase activity | 2 | 11 | 0.001 | 0.037 | DE >4fold |
| *G. hirsutum* | Maternal bias | IPR007524 | Pectate lyase, N-terminal | 2 | 11 | 0.002 | 0.037 | DE >4fold |
| *G. hirsutum* | Maternal bias | GO:0051082 | unfolded protein binding | 3 | 56 | 0.002 | 0.037 | DE >4fold |
| *G. hirsutum* | Maternal bias | IPR027413 | GroEL-like equatorial domain superfamily | 2 | 15 | 0.003 | 0.037 | DE >4fold |
| *G. hirsutum* | Maternal bias | IPR027409 | GroEL-like apical domain superfamily | 2 | 17 | 0.004 | 0.037 | DE >4fold |
| *G. hirsutum* | Maternal bias | IPR002423 | Chaperonin Cpn60/TCP-1 family | 2 | 18 | 0.004 | 0.037 | DE >4fold |
| *G. hirsutum* | Maternal bias | IPR008480 | Protein of unknown function DUF761, plant | 2 | 23 | 0.006 | 0.037 | DE >4fold |
| *G. hirsutum* | Maternal bias | IPR008972 | Cupredoxin | 3 | 76 | 0.006 | 0.037 | DE >4fold |
| *G. hirsutum* | Maternal bias | IPR025836 | Zinc knuckle CX2CX4HX4C | 2 | 24 | 0.007 | 0.037 | DE >4fold |
| *G. hirsutum* | Maternal bias | GO:0005524 | ATP binding | 12 | 1264 | 0.007 | 0.037 | DE >4fold |
| *G. hirsutum* | Maternal bias | IPR002022 | Pectate lyase | 2 | 26 | 0.008 | 0.037 | DE >4fold |
| *G. hirsutum* | Maternal bias | IPR018082 | AmbAllergen | 2 | 26 | 0.008 | 0.037 | DE >4fold |
| *G. hirsutum* | Maternal bias | IPR017451 | F-box associated interaction domain | 2 | 27 | 0.008 | 0.037 | DE >4fold |
| *G. hirsutum* | Maternal bias | GO:0006457 | protein folding | 3 | 92 | 0.008 | 0.037 | DE >4fold |
| *G. hirsutum* | Maternal bias | IPR019821 | Kinesin motor domain, conserved site | 2 | 28 | 0.009 | 0.037 | DE >4fold |
| *G. hirsutum* | Maternal bias | IPR025558 | Domain of unknown function DUF4283 | 2 | 28 | 0.009 | 0.037 | DE >4fold |
| *G. hirsutum* | Maternal bias | GO:0005871 | kinesin complex | 2 | 41 | 0.01 | 0.037 | DE >4fold |
| *G. hirsutum* | Maternal bias | GO:0003777 | microtubule motor activity | 2 | 44 | 0.016 | 0.037 | DE >4fold |
| *G. hirsutum* | Maternal bias | IPR001752 | Kinesin motor domain | 2 | 41 | 0.017 | 0.037 | DE >4fold |
| *G. hirsutum* | Maternal bias | IPR011707 | Multicopper oxidase, type 3 | 2 | 41 | 0.017 | 0.037 | DE >4fold |
| *G. hirsutum* | Maternal bias | IPR027640 | Kinesin-like protein | 2 | 41 | 0.017 | 0.037 | DE >4fold |
| *G. hirsutum* | Maternal bias | GO:0007018 | microtubule-based movement | 2 | 44 | 0.018 | 0.037 | DE >4fold |
| *G. hirsutum* | Maternal bias | IPR001117 | Multicopper oxidase, type 1 | 2 | 42 | 0.018 | 0.037 | DE >4fold |
| *G. hirsutum* | Maternal bias | IPR013763 | Cyclin-like | 2 | 43 | 0.019 | 0.037 | DE >4fold |
| *G. hirsutum* | Maternal bias | GO:0005507 | copper ion binding | 2 | 52 | 0.022 | 0.037 | DE >4fold |
| *G. hirsutum* | Maternal bias | GO:0008017 | microtubule binding | 2 | 53 | 0.023 | 0.037 | DE >4fold |
| *G. hirsutum* | Maternal bias | GO:0006950 | response to stress | 2 | 52 | 0.024 | 0.037 | DE >4fold |
| *G. hirsutum* | Maternal bias | GO:0004672 | protein kinase activity | 7 | 754 | 0.041 | 0.044 | DE >4fold |
| *G. hirsutum* | Maternal bias | GO:0006468 | protein phosphorylation | 7 | 754 | 0.047 | 0.047 | DE >4fold |
| *G. barbadense* | Paternal bias | IPR012340 | Nucleic acid-binding, OB-fold | 7 | 62 | 0.001 | 0.049 | any DE |
| *G. barbadense* | Paternal bias | IPR003851 | Zinc finger, Dof-type | 5 | 35 | 0.002 | 0.049 | any DE |
| *G. barbadense* | Paternal bias | IPR004853 | Sugar phosphate transporter domain | 4 | 29 | 0.006 | 0.049 | any DE |
| *G. barbadense* | Paternal bias | IPR023404 | Radical SAM, alpha/beta horseshoe | 2 | 4 | 0.007 | 0.049 | any DE |
| *G. barbadense* | Paternal bias | GO:0016620 | oxidoreductase activity, acting on the aldehyde or oxo group of donors, NAD or NADP as acceptor | 3 | 16 | 0.01 | 0.049 | any DE |
| *G. barbadense* | Paternal bias | IPR012394 | Aldehyde dehydrogenase NAD(P)-dependent | 2 | 5 | 0.01 | 0.049 | any DE |
| *G. barbadense* | Paternal bias | IPR029979 | Protein ESKIMO 1 | 2 | 5 | 0.01 | 0.049 | any DE |
| *G. barbadense* | Paternal bias | GO:0006081 | cellular aldehyde metabolic process | 2 | 5 | 0.011 | 0.049 | any DE |
| *G. barbadense* | Paternal bias | GO:0015780 | nucleotide-sugar transmembrane transport | 2 | 5 | 0.011 | 0.049 | any DE |
| *G. barbadense* | Paternal bias | GO:0050826 | response to freezing | 2 | 5 | 0.011 | 0.049 | any DE |
| *G. barbadense* | Paternal bias | IPR006638 | Elp3/MiaB/NifB | 2 | 6 | 0.013 | 0.049 | any DE |
| *G. barbadense* | Paternal bias | IPR016162 | Aldehyde dehydrogenase, N-terminal | 2 | 8 | 0.02 | 0.049 | any DE |
| *G. barbadense* | Paternal bias | GO:0006281 | DNA repair | 5 | 62 | 0.023 | 0.049 | any DE |
| *G. barbadense* | Paternal bias | IPR007197 | Radical SAM | 2 | 9 | 0.025 | 0.049 | any DE |
| *G. barbadense* | Paternal bias | IPR015590 | Aldehyde dehydrogenase domain | 2 | 9 | 0.025 | 0.049 | any DE |
| *G. barbadense* | Paternal bias | IPR016161 | Aldehyde/histidinol dehydrogenase | 2 | 9 | 0.025 | 0.049 | any DE |
| *G. barbadense* | Paternal bias | IPR016163 | Aldehyde dehydrogenase, C-terminal | 2 | 9 | 0.025 | 0.049 | any DE |
| *G. barbadense* | Paternal bias | IPR006689 | Small GTPase superfamily, ARF/SAR type | 3 | 25 | 0.025 | 0.049 | any DE |
| *G. barbadense* | Paternal bias | GO:0003684 | damaged DNA binding | 2 | 10 | 0.032 | 0.049 | any DE |
| *G. barbadense* | Paternal bias | IPR004146 | DC1 | 3 | 28 | 0.032 | 0.049 | any DE |
| *G. barbadense* | Paternal bias | GO:0006289 | nucleotide-excision repair | 2 | 11 | 0.038 | 0.049 | any DE |
| *G. barbadense* | Paternal bias | IPR012946 | X8 domain | 3 | 31 | 0.041 | 0.049 | any DE |
| *G. barbadense* | Paternal bias | IPR006068 | Cation-transporting P-type ATPase, C-terminal | 2 | 13 | 0.044 | 0.049 | any DE |
| *G. barbadense* | Paternal bias | IPR004000 | Actin family | 2 | 14 | 0.05 | 0.05 | any DE |
| *G. barbadense* | Maternal bias | IPR014001 | Helicase superfamily 1/2, ATP-binding domain | 5 | 61 | 0.003 | 0.049 | any DE |
| *G. barbadense* | Maternal bias | IPR001650 | Helicase, C-terminal | 5 | 62 | 0.003 | 0.049 | any DE |
| * *G. barbadense* | Maternal bias | GO:0009523 | photosystem II | 3 | 26 | 0.007 | 0.049 | any DE |
| *G. barbadense* | Maternal bias | IPR013922 | Cyclin PHO80-like | 2 | 7 | 0.008 | 0.049 | any DE |
| *G. barbadense* | Maternal bias | IPR014014 | RNA helicase, DEAD-box type, Q motif | 3 | 24 | 0.008 | 0.049 | any DE |
| *G. barbadense* | Maternal bias | GO:0000079 | regulation of cyclin-dependent protein serine/threonine kinase activity | 2 | 8 | 0.011 | 0.049 | any DE |
| *G. barbadense* | Maternal bias | GO:0019901 | protein kinase binding | 2 | 8 | 0.011 | 0.049 | any DE |
| *G. barbadense* | Maternal bias | IPR000198 | Rho GTPase-activating protein domain | 2 | 9 | 0.011 | 0.049 | any DE |
| * *G. barbadense* | Maternal bias | IPR002683 | PsbP, C-terminal | 2 | 9 | 0.011 | 0.049 | any DE |
| *G. barbadense* | Maternal bias | IPR008936 | Rho GTPase activation protein | 2 | 9 | 0.011 | 0.049 | any DE |
| * *G. barbadense* | Maternal bias | GO:0009654 | photosystem II oxygen evolving complex | 2 | 12 | 0.015 | 0.049 | any DE |
| *G. barbadense* | Maternal bias | GO:0019898 | extrinsic component of membrane | 2 | 12 | 0.015 | 0.049 | any DE |
| *G. barbadense* | Maternal bias | IPR002173 | Carbohydrate/purine kinase, PfkB, conserved site | 2 | 11 | 0.016 | 0.049 | any DE |
| *G. barbadense* | Maternal bias | IPR016123 | MogI/PsbP, alpha/beta/alpha sandwich | 2 | 11 | 0.016 | 0.049 | any DE |
| *G. barbadense* | Maternal bias | IPR000629 | ATP-dependent RNA helicase DEAD-box, conserved site | 2 | 13 | 0.021 | 0.049 | any DE |
| *G. barbadense* | Maternal bias | GO:0003950 | NAD+ ADP-ribosyltransferase activity | 2 | 12 | 0.022 | 0.049 | any DE |
| *G. barbadense* | Maternal bias | IPR027806 | Harbinger transposase-derived nuclease domain | 3 | 37 | 0.022 | 0.049 | any DE |
| *G. barbadense* | Maternal bias | IPR011545 | DEAD/DEAH box helicase domain | 3 | 39 | 0.026 | 0.049 | any DE |
| *G. barbadense* | Maternal bias | IPR011611 | Carbohydrate kinase PfkB | 2 | 15 | 0.027 | 0.049 | any DE |
| *G. barbadense* | Maternal bias | IPR013763 | Cyclin-like | 3 | 41 | 0.029 | 0.049 | any DE |
| *G. barbadense* | Maternal bias | IPR029056 | Ribokinase-like | 2 | 16 | 0.03 | 0.049 | any DE |
| *G. barbadense* | Maternal bias | IPR019794 | Peroxidase, active site | 3 | 42 | 0.031 | 0.049 | any DE |
| * *G. barbadense* | Maternal bias | GO:0015979 | photosynthesis | 3 | 40 | 0.032 | 0.049 | any DE |
| *G. barbadense* | Maternal bias | IPR024709 | Putative O-fucosyltransferase, plant | 2 | 18 | 0.036 | 0.049 | any DE |
| *G. barbadense* | Maternal bias | IPR025525 | hAT-like transposase, RNase-H fold | 2 | 18 | 0.036 | 0.049 | any DE |
| *G. barbadense* | Maternal bias | IPR021720 | Malectin domain | 2 | 19 | 0.04 | 0.049 | any DE |
| *G. barbadense* | Maternal bias | GO:0046983 | protein dimerization activity | 8 | 225 | 0.043 | 0.049 | any DE |
| *G. barbadense* | Maternal bias | IPR000823 | Plant peroxidase | 3 | 49 | 0.044 | 0.049 | any DE |
| *G. barbadense* | Paternal bias | GO:0006281 | DNA repair | 5 | 62 | 0 | 0.014 | DE >4fold |
| *G. barbadense* | Paternal bias | IPR003851 | Zinc finger, Dof-type | 3 | 35 | 0.003 | 0.036 | DE >4fold |
| *G. barbadense* | Paternal bias | GO:0003684 | damaged DNA binding | 2 | 10 | 0.004 | 0.036 | DE >4fold |
| *G. barbadense* | Paternal bias | GO:0006289 | nucleotide-excision repair | 2 | 11 | 0.007 | 0.036 | DE >4fold |
| *G. barbadense* | Paternal bias | IPR006626 | Parallel beta-helix repeat | 2 | 28 | 0.021 | 0.036 | DE >4fold |
| *G. barbadense* | Paternal bias | IPR008928 | Six-hairpin glycosidase superfamily | 2 | 29 | 0.022 | 0.036 | DE >4fold |
| *G. barbadense* | Paternal bias | IPR000743 | Glycoside hydrolase, family 28 | 2 | 35 | 0.03 | 0.037 | DE >4fold |
| *G. barbadense* | Paternal bias | GO:0004650 | polygalacturonase activity | 2 | 35 | 0.036 | 0.04 | DE >4fold |
| *G. barbadense* | Maternal bias | IPR013922 | Cyclin PHO80-like | 2 | 7 | 0 | 0.014 | DE >4fold |
| *G. barbadense* | Maternal bias | IPR013763 | Cyclin-like | 3 | 41 | 0 | 0.014 | DE >4fold |
| *G. barbadense* | Maternal bias | GO:0000079 | regulation of cyclin-dependent protein serine/threonine kinase activity | 2 | 8 | 0 | 0.014 | DE >4fold |
| *G. barbadense* | Maternal bias | GO:0019901 | protein kinase binding | 2 | 8 | 0 | 0.014 | DE >4fold |
| *G. barbadense* | Maternal bias | IPR027806 | Harbinger transposase-derived nuclease domain | 2 | 37 | 0.006 | 0.036 | DE >4fold |

